# G-matrix stability in clinally diverging populations of an annual weed

**DOI:** 10.1101/2022.03.08.483458

**Authors:** Georgia Alexandria Henry, John R Stinchcombe

## Abstract

How phenotypic and genetic divergence among populations is influenced by the genetic architecture of those traits, and how microevolutionary changes in turn affect the within-population patterns of genetic variation, are of major interest to evolutionary biology. Work on *Ipomoea hederacea*, an annual vine, has found genetic clines in the means of a suite of ecologically important traits, including flowering time, growth rate, seed mass, and corolla width. Here we investigate the genetic (co)variances of these clinally varying traits in two northern range-edge and two central populations of *Ipomoea hederacea* to evaluate the influence of the genetic architecture on divergence across the range. We find 1) limited evidence for clear differentiation between Northern and Southern populations in the structure of **G**, suggesting overall stability of **G** across the range despite mean trait divergence and 2) that the axes of greatest variation (g_max_) were unaligned with the axis of greatest multivariate divergence. Together these results indicate the role of the quantitative genetic architecture in constraining evolutionary response and divergence among populations across the geographic range.

## Introduction

A major goal in evolutionary biology is to understand the relationship between genetic variation within populations and phenotypic divergence between populations (Antonovics 1976, Endler 1977, Walsh & Blows 2009). Range-edge populations provide an interesting setting in which to investigate this relationship. Populations existing on the margin of species’ ranges are expected to have smaller population sizes (Brown et al. 1995), which are more susceptible to drift, and limited gene flow compared to more central populations (Sexton et al. 2009). Environmental gradients will often additionally create different selective pressures across the geographic range, favouring phenotypic and genetic divergence among populations (Endler 1977, Slatkin 1978, Barton 2001). Here we examine the divergence and evolutionary potential of four populations of an annual plant, *Ipomoea hederacea*, sampled from the center and northern edge of its range with respect to five clinally varying traits.

The evolutionary response to selection is dependent not only on local selection, but on the underlying genetic architecture of the traits undergoing selection. The ability to adapt to the local environment depends on both standing genetic variation and covariation, represented as a genetic covariance matrix, **G** (Antonovics 1976, Lande 1979, Lande & Arnold 1983, Falconer & Mackay 1996, Agrawal & Stinchcombe 2009). **G** shapes evolutionary responses to selection, as the relationships among traits lead to correlated responses (Lande 1979, Lande & Arnold 1983). Divergence among populations in response to local selection may thus be constrained or facilitated by trait (co)variances. The multivariate trait combination with the greatest amount of genetic variation, dubbed the “genetic line of least resistance” by Schluter (1996) has been shown to influence macroevolutionary responses by deflecting the response to selection toward trait combinations with the most genetic variation (Schluter 1996).

Exactly how G-matrices will change with divergence in trait means across latitudinal ranges is not clear, although there are numerous reasons to predict that they will. For example, while some work suggests that G-matrices are expected to exhibit stability over various evolutionary timescales (Lande 1979, Schluter 1996, Arnold et al. 2008), this assumes genes underlying traits are from an infinite-sites model with a gaussian distribution (Lande 1979). **G** may be unstable through time if the distribution of the alleles underlying traits is skewed (Barton & Turelli 1987, Turelli 1988). Additionally, **G** among populations may differ if a population is small and subject to substantial genetic drift or inbreeding (Lande 1979, Phillips et al. 2001, Jones et al. 2004, Doroszuk et al. 2008), has experienced recent population bottlenecks (Shaw et al. 1995, Roff 2000), or is under strong selection due to novel conditions (Turelli 1988, Roff 2000, Conner et al. 2011, Uesugi et al. 2017), which are all features often associated with range-edge populations. Recent theory by Chantepie and Chevin (2020) suggests that in finite populations subject to drift, genetic correlations will more closely reflect mutational (co)variances as effective population sizes become smaller. Chantepie and Chevin (2020) note that colonizing populations (which may be similar to range-edge populations) often experience reduced Ne and strong selection, and thus **G** will be more reflective of mutational covariances, and stronger genetic constraints. Taken together, we might predict less overall genetic variance (Johnson & Barton 1995), or different patterns of genetic covariance, which may in turn lead to evolutionary constraint in range-edge populations.

There are, however, some scenarios in which **G** might be predicted to be similar among populations. If range-edge populations frequently experience swamping gene flow from the center of the range (Kirkpatrick & Barton 1997; Garcia-Ramos & Kirkpatrick 1997), their **G** might show few differences compared to core populations, with populations diverging along the homogenized axis of maximal (co)variation (Guillaume & Whitlock 2007). Some theory also suggests that dispersal into peripheral populations can enhance genetic variance (counteracting the effects of drift), facilitating adaptation to range-edge conditions (Polechova 2015, Polechova 2018).

Empirical assessment of divergence among population G-matrices in natural populations has led to mixed results. Some studies find that **G** is fairly stable (Roff & Mousseau 1999, Puentes et al. 2016, Delahaie et al. 2017, McGoey & Stinchcombe 2021) with most divergence among populations occurring along g_max_ (Silva e Costa et al. 2020, Royaute et al. 2020). In Australian populations of *Drosophila melanogaster* sampled along a broad environmental gradient, Hangartner et al. (2019) estimated G-matrices for size, desiccation, and thermal traits. They found no differentiation among population G-matrices, demonstrating the robustness of **G,** despite diverging local selection. In contrast, Hine et al. (2009) evaluated the divergence in the G-matrices for cuticular hydrocarbons of nine *Drosophila serrata* populations along the same gradient, and found significant divergence among **G**s. Wood and Brodie (2015) showed that environmental effects on **G** (e.g., due to novel habitats) were often as large, or larger, than population differentiation, suggesting that environmental sensitivity in the expression of **G** can have effects as large as the accumulated effects of drift, selection, and mutation between diverging populations. Similarly, some studies have found remarkable shifts in **G** through time and/or across environments (Cano et al. 2004, Doroszuk et al. 2008, Bjorklund et al. 2013).

Collectively, this diverse body of theory and empiricism suggests numerous reasons why **G** from range-edge and populations from the interior of a species range should differ. Under a wide range of biologically plausible assumptions about genetic architecture, inbreeding, drift, effective population size, bottlenecks, dispersal, and strong selection—all of which are expected to be relevant to range-edge populations and those adapting to a gradient—lead to the prediction of differences in G. However, alternative theoretical assumptions about genetic architecture or divergence could lead to predictions of no or little change in **G**. Past empirical work likewise offers contrasting guidance: **G** might show minimal or alternatively strong changes due to temporal or environmental effects. Consequently, whether range-edge populations that have diverged along a gradient show altered **G –** and thus potentially altered evolutionary potential and constraint **–** remains an unanswered empirical question.

*Ipomoea hederacea,* an annual plant which grows across the eastern U.S.A. from Pennsylvania to Florida, has latitudinal genetic clines in a variety of quantitative traits (Klingaman and Oliver 1996, Stock et al. 2014) and leaf shape (Bright-Emlen 1998, Campitelli & Stinchcombe 2013), a Mendelian trait. These clines do not appear to be due to drift, but rather a response to selection, as no latitudinal patterns in neutral loci have been found (Campitelli & Stinchcombe 2013; Campitelli & Stinchcombe 2014). Previous work by Stock and colleagues (2014) focused on among-population patterns found evidence of significant clinal divergence.

Briefly, Stock et al. (2014) grew individuals in a common greenhouse environment from 20 populations, spanning the gradient sampled by Campitelli and Stinchcombe (2013) . They used selfed seeds from 10 maternal lines sampled per population and up to 8 individuals per matriline. They performed redundancy analysis using the matriline means of five focal traits (seed mass, early growth rate, flowering time, corolla width, and anther-stigma distance) regressed on latitude of origin. Flowering time, anther-stigma distance, and corolla width exhibited significant latitudinal clines, while seed mass and growth rate had marginally significant divergence. Stock et al. (2014) estimated the axis of greatest multivariate divergence, the mean trait combination that is most different between populations of *I. hederacea* across latitudes, which we use here alongside more deeply sampled populations to compare within-population trait organization to among-population trait divergence.

Our research aims were: 1) to determine whether Northern and Southern populations differed in the overall heritability and genetic variance of the focal traits, 2) evaluate how genetic variation within populations was aligned relative to the axis of multivariate genetic divergence, and 3) assess population level divergence of the G-matrices. To do so, we estimated trait heritability and genetic variance-covariance matrices of four populations of *Ipomoea hederacea*; collected from the northern range-edge and the core of the range. We expected the axis of trait divergence between populations to be biased toward the axis of greatest variation within populations, the genetic line of least resistance (Schluter 1996).

## Methods

### Propagation and trait measurements

We harvested seed from populations from the edge of agricultural fields in Pennsylvania (40.116611°N, 76.398889°W), Maryland (39.581861°N, 77.816861°W), Hoffman, North Carolina (35.074139°N, 79.556694°W) and Ellerbe, North Carolina (35.091722°N, 79.742722°W). We sampled seed pods haphazardly from vines 1-2 m from one another to minimize repeat sampling of maternal plants, which served as the founders of the matrilines we used in our experiment. We grew one seed from 50 randomly chosen matrilines per population in a common greenhouse environment and allowed these individuals to self-fertilize. One death prior to seed set resulted in 49 matrilines from Ellerbe, NC. We harvested selfed seeds from each maternal plant and grew ∼11 focal individuals (range: 2-11, mean = 10.7) from each line, resulting in a total sample of 2,133 plants. While our use of selfed progeny precludes estimating additive genetic variance, selection acts on the broad-sense variation rather than just the additive components of variation in mainly selfing species (Roughgarden 1979), making this design appropriate for *I. hederacea*.

We grew plants in cone-tainers (Stuewe & Sons, Oregon, USA) filled with Pro-Mix soil in a glasshouse from April-October 2018. All plants received a dilute fertilizer solution (N-P-K: 10-52-10) every 2 weeks. Day length was initially 16 hours at 28°C, reduced to 12 hours at 25℃ on day 32, and 8 hours at 22°C on day 38 to promote flowering.

Trait measurements followed Stock et al. (2014). Prior to planting, we weighed each seed and recorded individual mass. We counted leaves at 17 days after planting and again on the 26th or 27th day after planting; we estimated growth rate as the difference in leaf counts divided by the number of days, and as such it is in units of new leaves/day. We measured corolla width and anther-stigma distance on the first flower of each individual using digital calipers. We measured the distance from the lowest and highest anthers to the stigma and calculated the mean of the absolute distance to characterize anther-stigma distance.

### Data preparation

Four of the five focal traits, (seed mass, early growth rate, corolla width and anther- stigma distance) were standardized to mean = 0 and standard deviation = 1 using the grand mean and standard deviation across populations to allow for meaningful comparison of differences (Hine et al. 2009, Hansen & Houle 2008). We elected to use this standardization to eliminate differences in the scale and units of our traits (e.g., anther-stigma distance in mm, growth rate in leaves per day, and flowering time in days) and because mean standardization can be difficult to apply to traits like flowering time, which have an arbitrary rather than natural zero (Hansen & Houle 2008, Houle et al. 2011). During the experiment the shift from lengthening to shortening days interrupted the flowering schedule. The glasshouse in which the plants were grown was exposed to ambient light, and the change in artificial light used to induce flowering was largely overwhelmed by the natural photoperiod. For this reason, we transformed the number of days until the first flower using ordered quantile normalization (ORQ), which preserves the order of flowering but results in a gaussian distribution (Peterson and Cavanaugh 2019). The ORQ normalization results in a distribution centered on zero with a standard deviation of one, which is the same scale as all other traits.

### Univariate analysis

To determine broad-sense heritabilities of the traits we fit Bayesian generalized linear mixed-effect models for each trait across all populations. We estimated the models using *MCMCglmm* in R (Hadfield 2010) using the mixed model,

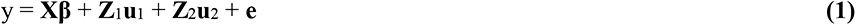

For each trait, y, we ran a separate model. In (1), **X**, **Z**_1_ and **Z**_2_ represent design matrices for the fixed effect of population, and random effects of matriline and greenhouse block, respectively. **β** and **u** are vectors of the related parameters. We included the environmental block to control for the effect of microenvironmental differences within the glasshouse. For seed mass, we used the greenhouse block of the maternal plant. Residual error is represented by **e.**

We tested six priors and summed the Deviance Information Criteria value for each of the five univariate models. We used the prior with the lowest summed DIC score, although due to the amount of data in the model, the prior was not very influential. The effective size of the samples was greater than 85% of the total number of samples and the autocorrelation was below 0.05 for all parameters (Hadfield 2010). To evaluate whether there was meaningful broad-sense heritability, we ran models within each population with and without the matriline variable and compared DIC values, per Puentes et al. (2016). We considered models where the matriline reduced the DIC score >2 to have a “significant” genetic component and thus H^2^ was significant. We constructed comparable models for validation, using REML, which are presented in the Supplemental Material. Heritability estimates were similar, with the greatest difference in Heritability of 5.2%, and an average absolute difference of 1.7%. The 95% Highest Posterior Density (HPD) intervals from Bayesian models overlapped REML heritability estimates in all cases, and as such Bayesian estimates are presented in the main text.

### Multivariate Analyses

We estimated the total genetic variance-covariance matrices separately for each population using Bayesian generalized linear mixed-effect models through *MCMCglmm* in R (Hadfield 2010). We again used the mixed model,

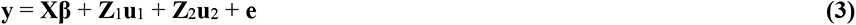

where **X**, **Z**_1_ and **Z**_2_ represent design matrices for the vectors of trait means (**β**), the total genetic effects (**u**_1_) and the greenhouse block effects (**u**_2_). The residual error is represented by the term **e**.

We used weakly informative inverse-Wishart priors for estimating the variance and covariances where the distribution was described by the variance (V) and the belief parameter (nu) and the default prior for the fixed effect estimates, *N*(0,10^8^). We tested a variety of priors to determine the robustness of estimates and selected the model with the best fitting prior, determined by the lowest average Deviation Information Criteria score to use for our analyses (see Supplementary Materials). The final prior for the residual variance was given by a diagonal matrix of 1’s with the degree of belief slightly above *n*-1, for *n* traits (V=diag(5), nu = 4.001).

The best prior for the random effects was a matrix of 1/6 the phenotypic variance-covariance matrix and a belief parameter slightly above *n*-1 (V = (**P**/6)×100, nu=4.001, where **P** is the phenotypic (co)variance matrix). All the response variables in the model were multiplied by a factor of 10, which improves the estimation of small variances (and is subsequently factored out of the genetic variance-covariance matrix). To account for this factor of 10 in the response variables, the Phenotypic variance-covariance matrix (**P**) was multiplied by a factor of 100 in the priors.

We ran full models for 5,005,000 iterations, with a burn in of 5,000 and a thinning interval of 1,000, resulting in 5000 iterations sampled from the posterior distribution. As with our univariate models, the effective size of the samples were greater than 85% of the total number of samples and all parameters had <0.05 autocorrelation (Hadfield 2010).

### Null Models

For some analyses a null model was necessary, as MCMC methods can only estimate variance greater than zero (matrices must be positive definite). We constructed randomized G-matrices where the values only reflected sampling error as a point of comparison to our actual estimates. Following Walter et al. (2018) and McGoey and Stinchcombe (2021), we randomly assigned individuals to matrilines within populations, without replacement, resulting in G-matrices that are only due to sampling error, as any phenotypic similarity among individuals is only due to random sampling, rather than true genetic ancestry. Our design includes significant fixed effects, which Morrissey et al. (2019) note will make these null distributions larger than they should be, which should make our approach conservative. We also note that because each randomized G-matrix only reflects sampling variation, the expectation of a large number of G-matrix comparisons should be near 0 (as each population only reflects sampling variation), but the distribution of these comparisons reflects the range of potential outcomes one can obtain if each **G** only reflects sampling, making it a sensible null model. The within-population randomization approach also allows us to compare whether each estimated **G** differs from expectations due to sampling (Sztepanacz and Blows 2017).

We fed each randomized “population” into the same models as the observed data, run for 20,000 iterations with a burn in of 5,000 and a thinning interval of 100. We iterated the randomization and model estimates 1000 times. We combined the final posterior sample of each of the randomized models within populations to be used as a composite posterior. We assessed chains visually for stability and with Gelman and Rubin convergence diagnostic criteria.

### G-matrix comparisons

We used several metrics to compare G-matrices, each of which has different strengths and weaknesses (Krzanowski 1979, Kirkpatrick 2009, Hine et al 2009, Aguirre et al. 2014, Teplitsky et al. 2014, Puentes et al. 2016, Walter et al. 2018). In general, our goal was to use a variety of metrics to compare G-matrix size, concordance between patterns of genetic variance and clinal divergence, and similarity in the responses to selection that each G-matrix would produce.

### G-matrix size

We calculated the trace of the G-matrices to determine the total amount of genetic variance in each population. The trace then describes the overall size of the G-matrices, or the total amount of quantitative genetic variation they contain, which may be smaller in populations affected by recent bottlenecks or strong selection as genetic variation is lost or eroded. We tested our hypotheses using the mean of the posterior distributions and the 95% HPD intervals, to examine whether there were differences in quantitative genetic variance. We also utilized the posterior distribution of **G** to retrospectively evaluate how different they would need to be to detect differences in the trace. To do so, we multiplied the posterior distribution of the Northern populations by successive fractions to determine how much shrinkage would be required for the 95% HPD interval of a Northern population to not overlap that of the Southern populations. Our logic was that if only minor reductions in **G** were required (e.g., the fraction is 0.95), any conclusion of similarity would be relatively weak, and suggest a combination of low power, and sampling or estimation error. In contrast, if detecting a difference between Northern and Southern **G** required dramatic shrinkage of the matrix (e.g., the fraction is 0.25), it would suggest that any conclusion about trace similarity was robust.

### Genetic variation in the direction of maximum clinal divergence

To evaluate the alignment between the axis of greatest genetic variation (g_max_) of each population, and between g_max_ and the vector of greatest multivariate divergence (Cline_max_ from Stock et al. 2014) we calculated the correlation coefficient between these vectors. We used random unit vectors as a null comparison. We note that Stock et al. (2014) used and presented results from mean-standardized data; we re-analyzed their data to estimate the vector of greatest clinal divergence using standard deviation standardized data, to match our use of standard deviation-standardized data from the greenhouse experiment.

We projected the vector of maximum clinal divergence through each G-matrix (using b^T^**G**b, where b is the vector of clinal divergence; see Lin and Allaire 1977) to calculate the genetic variance in direction of clinal divergence. To estimate the proportion of genetic variance along the direction of maximum clinal divergence, we estimate b^T^**G**b/λ_1_, where λ_1_ is the first eigenvalue of **G**, representing the vector of maximum genetic variance in each population. Given that the G-matrices had different shapes, the maximum amount of genetic variation in any direction differed between the populations. To allow for fair comparison, we standardized by the proportion of variance associated with g_max_, the first eigenvector of **G**, in each population.

### Random skewers

We performed a random skewers analysis (Cheverud & Marriog 2007, Aguirre et al. 2014) following Aguirre et al. 2014. Briefly, 1000 randomly generated normal vectors were projected through each population’s **G**. We compared the response from each projection between populations to determine which random vectors which produced response differences. We then calculate the variance-covariance matrix of the skewers which result in response differences between populations and performed eigenanalysis to generate the **R** matrix. The **R** matrix describes the trait space where differences in the response to skewers exist between populations, with the first eigenvector describing the axis with the greatest differences between populations.

### Krzanowski’s subspace analysis

Krzanowski’s subspace analysis provides a method of evaluating the similarity in geometry of multiple variance-covariance matrices. The approach works by examining whether the subspaces containing most of the genetic variation (e.g., the leading PCs) in each population are shared, or in common. We first determine the shared space (**H**) of all populations, from a subset of the leading PCs (eigenvectors) of each population, using:

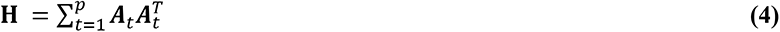

where superscript T indicates transposition, and a subset *t* of PCs of each G-matrix are included in **A**, and summation is to *p* populations (*p* = 4 in our case; Krzanowski 1979; Aguirre et al. 2014). As per Aguirre et al. 2014, the number of eigenvectors used for each population were those which explained >90% of the total variation to account for differences in the shape of the population G-matrices. The eigenvalues of **H** can range from 0 to *p,* which indicates common sub-spaces to the matrices being compared (i.e., the leading PCs describe similar directions of multivariate space). If the eigenvalues of **H** are less than *p* (4 in our case), this indicates that genetic variation described by that eigenvector differs among the populations. To evaluate which populations are most similar we determine the smallest angle between the mean posterior subspace of each population and the mean shared subspace **H**, as per Aguirre et al. 2014.

### Tensor analysis

Differences between matrices can be evaluated by calculating the 4^th^-order covariance tensor, describing the variances and covariances between matrices (Hine et al. 2009). We used the 4^th^-Order Genetic Covariance Tensor analysis developed by Hine et al. (2009), and outlined by Aguirre et al. (2014), and Walter et al. (2018). We refer interested readers to these papers for full details (especially Fig. 2 of Walter et al. 2018), and only provide a brief overview here. The fourth-order tensor of the matrices describes the variance and covariances among the population G-matrices. The tensor can be represented as a symmetric (n(n+1)/2) matrix (**S**) (Basser & Pajevic 2007), where n is the number of traits, with elements describing the variances and covariances of genetic variances and covariances among the four populations. The matrix **S** can then be subject to eigenanalysis followed by scaling and rearrangement, such that the eigenvalues and eigentensors describe how the G-matrices vary. Eigentensors with larger eigenvalues (which explain a larger percentage of variance) describe dimensions of greater variation in the G-matrices; the eigentensors can themselves be subjected eigenanalysis, to determine the linear combination of traits (eigenvectors) that lead to the difference in G-matrices captured by the eigentensors. The covariance tensor analysis can thus be used to evaluate axes of differences among the genetic variance-covariance structure of the populations.

## Results

### Univariate Analyses

We estimated broad-sense heritability of each trait within populations using a Bayesian model (Table S1, Figure S1), and confirmed results with REML (Table S2). All traits had heritability greater than zero in at least 3 of the 4 populations (Table S1). The Pennsylvania population lacked significant heritability for anther-stigma distance (Table S1). The mean of all the heritability estimates was 0.213 (ranging from 0.011 to 0.535). There was no significant difference between Northern and Southern populations, although Northern populations tended to have lower heritability estimates than Southern populations (Table S1).

### G-matrix size

We present population G-matrices, 95% HPD intervals, and null expectations due to sampling variation in the Supplemental Material (Table S3, Table S4), and focus here on comparisons of the matrices. The G-matrices all contain variances with HPD intervals that do not overlap the 95% HPD intervals of the null G-matrices, although some traits in some populations did have overlapping 95% HPD intervals, indicating that the variation may be very small (Table S3). Two traits, corolla width and anther-stigma distance, have variation indistinguishable from sampling variation in all populations except Hoffman (Table S3); we nonetheless retained these traits in our analyses to allow comparisons of **G** to divergence in trait means, which requires these traits.

We found that Hoffman, NC had significantly greater total genetic variation than either Northern population, estimated as the trace of the G-matrices. Hoffman had approximately 57% greater genetic variation than populations from Ellerbe, NC, 84% greater variation than Maryland, and 142% greater variation than Pennsylvania. The 95% HPD intervals of Hoffman did not overlap with either Northern population, but did overlap slightly with the other Southern population, Ellerbe, NC.

We next manipulated the posterior distribution to determine the proportional change required to eliminate the overlap of the 95% HPD intervals between Northern and Southern populations (taking advantage of the fact that posterior distributions remain valid for operations performed on them: Wilson et al. 2010). As the trend was for smaller trace values in the north, which we expect in recently bottlenecked populations or those eroded by selection, we calculated the necessary reductions in the trace for Northern populations. Hoffman, NC was already detectably different from the Northern populations, and so Ellerbe was the contrast for this assessment. A reduction of 15% of the trace from the estimated trace of Pennsylvania and 50% reduction of Maryland were sufficient to allow for differentiation between those populations and Ellerbe. These analyses suggest that the Pennsylvania population may in fact have a reduced trace but that estimation and sampling error, and power, preclude us detecting it. In contrast, the trace for the Maryland population appears to be robustly similar to Southern populations.

### Genetic variation in the direction of maximum clinal divergence

Overall, the proportion of genetic variation in the direction of maximum clinal divergence was small, roughly on par with the third eigenvalues of each population, and there were no significant differences between any populations (Table 2). To fairly compare between populations, which differ in their total amount of genetic variation (Table 1) and the variation along g_max_ (Table S5), we standardized the genetic variance in the direction of clinal divergence by the genetic variance associated with g_max_, so the values are relative to the linear combination of traits with the maximum possible genetic variance in each population. The standardized projections of the Northern populations were slightly larger than the Southern populations, but there were no significant differences between them. In general, genetic variance in the direction of clinal variation was ∼30-50% of the genetic variance associated with g_max_.

**Table 1:** The total amount of genetic variation in each population estimated as the trace of **G**. Low and high values demarcate the 95% highest posterior density interval (HPD). The trace of the randomized null G-matrices are included for comparison.

| Population | Trace of G | Lower HPD | Upper HPD | Random Trace | L. HPD | U. HPD |
| --- | --- | --- | --- | --- | --- | --- |
| Pennsylvania | 0.708 | 0.524 | 0.896 | 0.137 | 0.078 | 0.195 |
| Maryland | 0.930 | 0.689 | 1.178 | 0.138 | 0.085 | 0.209 |
| Hoffman, NC | 1.716 | 1.311 | 2.181 | 0.153 | 0.093 | 0.232 |
| Ellerbe, NC | 1.091 | 0.799 | 1.386 | 0.148 | 0.092 | 0.216 |

**Table 2:** Genetic variance in the direction of maximum clinal divergence for each population. In the standardized column (Variance (std)), the variances are standardized by the eigenvalue of the leading eigenvector of each population, to express this relative to the direction of greatest genetic variation. Lower and higher HPD values demarcate the 95% highest posterior density interval.

| Population | Variance | Lower HPD | Higher HPD | Variance (std) | Lower HPD | Higher HPD |
| --- | --- | --- | --- | --- | --- | --- |
| Pennsylvania | 0.155 | 0.076 | 0.241 | 0.505 | 0.258 | 0.782 |
| Maryland | 0.212 | 0.111 | 0.326 | 0.492 | 0.274 | 0.721 |
| Hoffman, NC | 0.247 | 0.130 | 0.380 | 0.321 | 0.139 | 0.523 |
| Ellerbe, NC | 0.283 | 0.151 | 0.436 | 0.591 | 0.319 | 0.869 |

The correlation between g_max_ from each of the G-matrices and direction of maximum clinal divergence is not significantly different from the expected correlation between g_max_ and a random vector (Figure 1). Although we might expect the direction of maximum clinal divergence to be aligned with directions of genetic variation, this does not appear to be the case for any population. The correlation between PC2 through PC5 of the population G-matrices and direction of maximum clinal divergence followed the same pattern (Figure S2).

**Figure 1:**
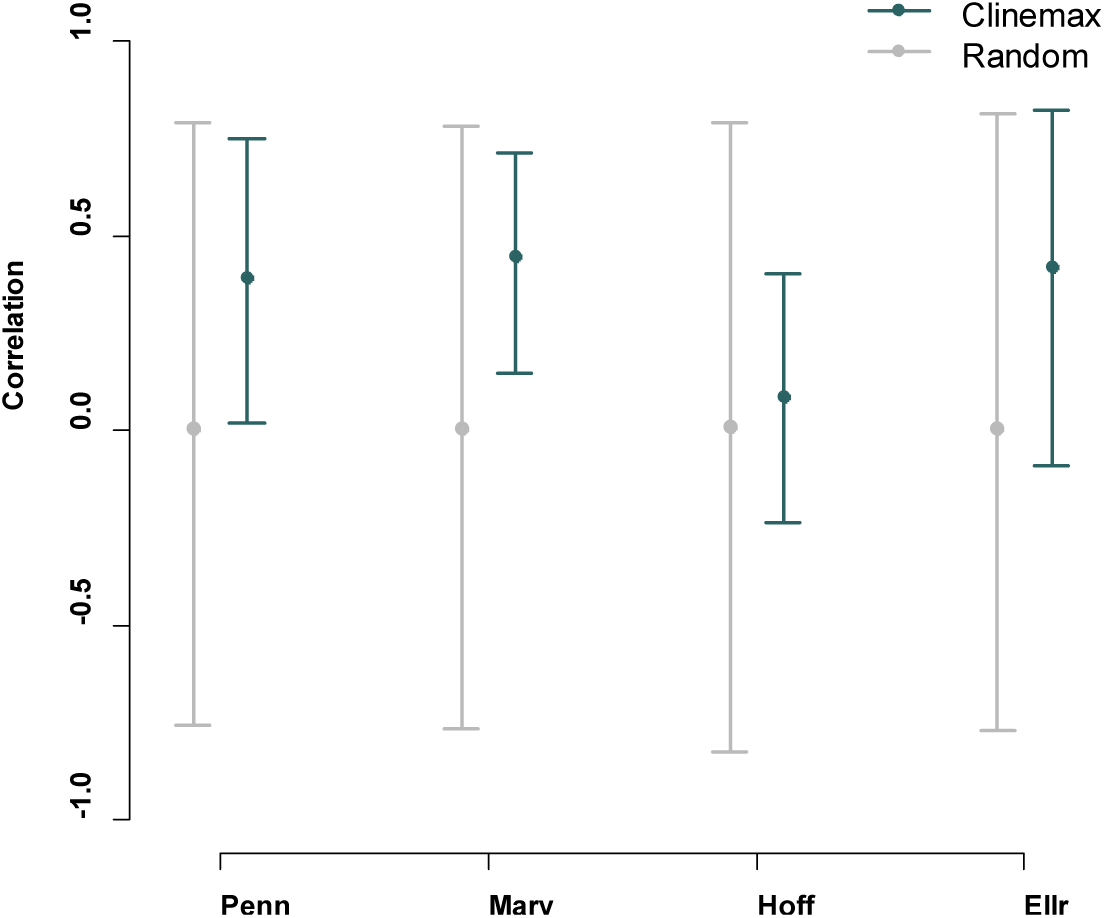
The correlation between the g_max_ of each population G-matrices and the vector of greatest multivariate clinal divergence (Observed, in blue) compared to the correlation between the g_max_ of each population G-matrices and randomized vectors (Randomized, in, in gray). Error bars represent the 95% HPD intervals. “Penn” is Pennsylvania, “Mary” is Maryland, “Hoff” is Hoffman, NC and “Ellr” is Ellerbe, NC.

### Random skewers

The considerable fraction of the random vectors produced some difference in response between populations, with 350 of 1000 resulting in responses where at least one pairwise comparison had 95% HPD intervals which did not overlap. Following Aguirre et al. (2014), we took this subset of vectors which resulted in response differences and created a variance-covariance matrix of the vectors. We then performed eigenanalysis to evaluate the trait space which describes differences among populations. The highest trait loading of the first eigenvector is corolla width, followed by seed mass and flowering time (table 3). These differences in response are largely attributable to Hoffman, NC. Projecting the eigenvectors of the R-matrix through each population’s G-matrix, we found that Hoffman had significantly more genetic variation along this eigenvector than the other three populations (Figure 2). In other words, the response difference we detected with random skewers appears to be due to the Hoffman population having more genetic variation for a linear combination of traits made up primarily of corolla width, flowering time and seed mass, with lesser contributions of growth rate and anther stigma distance, than the other three populations.

**Table 3:** Eigenanalysis of the R-matrix from the Random Skewers analysis which resulted in difference in response when projected through populations. The R-matrix summarized the trait combinations which resulted in differences to the projected response. Each eigenvector (RSe1 through RSe5) describes an independent combination of traits which capture differences among populations. The eigenvalues describe the proportion of variation of each eigenvector within **R**, larger values indicate more of the variation in the skewers are explained.

| Trait | RSe1 | RSe2 | RSe3 | RSe4 | RSe5 |
| --- | --- | --- | --- | --- | --- |
| Seed mass | -0.471 | -0.153 | 0.402 | -0.404 | 0.656 |
| Growth | -0.041 | -0.045 | 0.811 | -0.069 | -0.578 |
| Flow time | -0.513 | -0.525 | -0.034 | 0.676 | -0.053 |
| Corolla | -0.655 | 0.158 | -0.388 | -0.429 | -0.459 |
| AS dist. | -0.292 | 0.821 | 0.172 | 0.436 | 0.145 |
| **Eigenvalues** | **0.419** | **0.230** | **0.177** | **0.106** | **0.068** |

**Figure 2:**
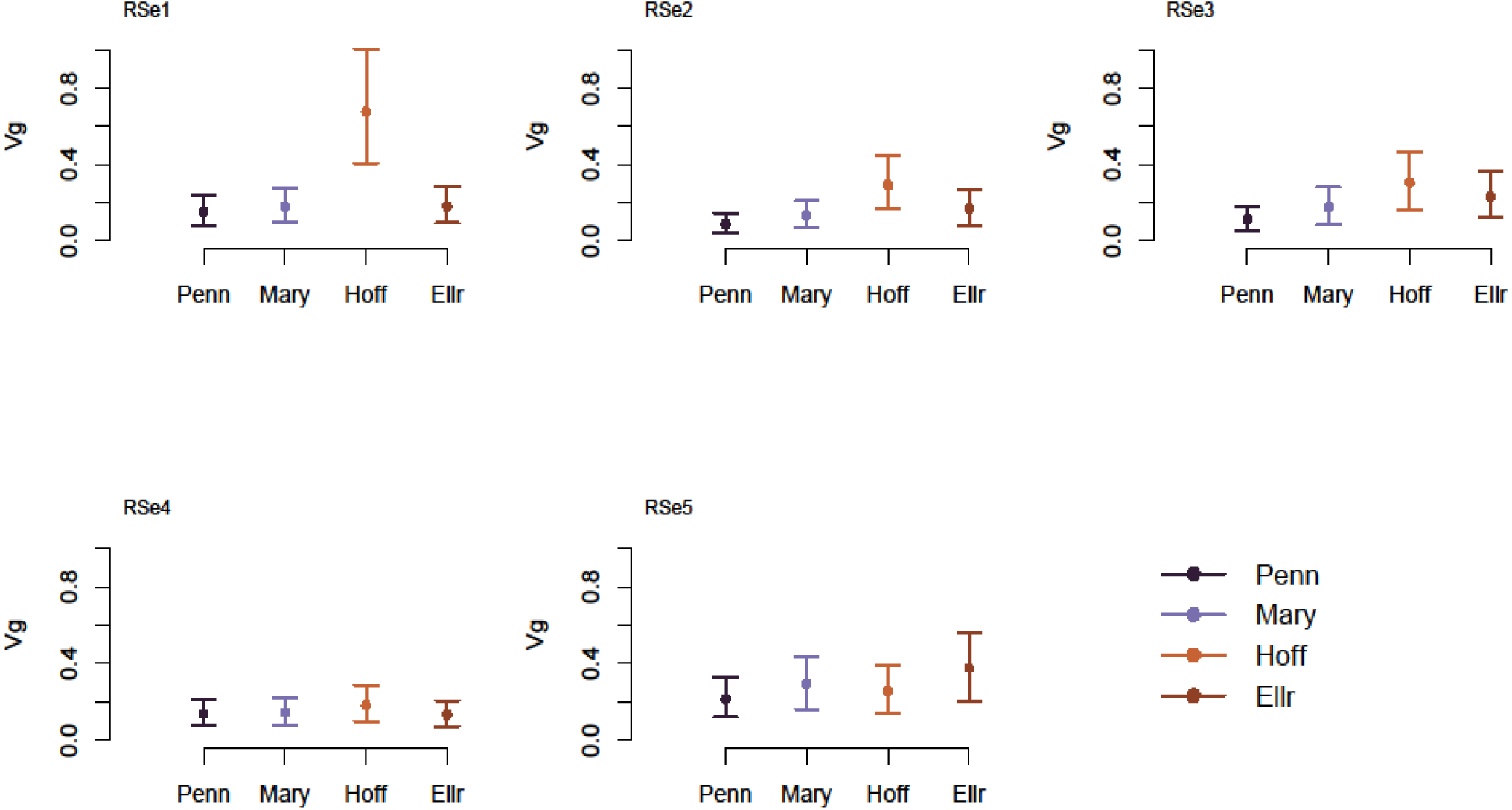
Genetic variance (Vg) in the direction of each eigenvector of the **R** matrix, where the **R** matrix is composed of vectors which resulted in response differences between populations.

**Figure 3:**
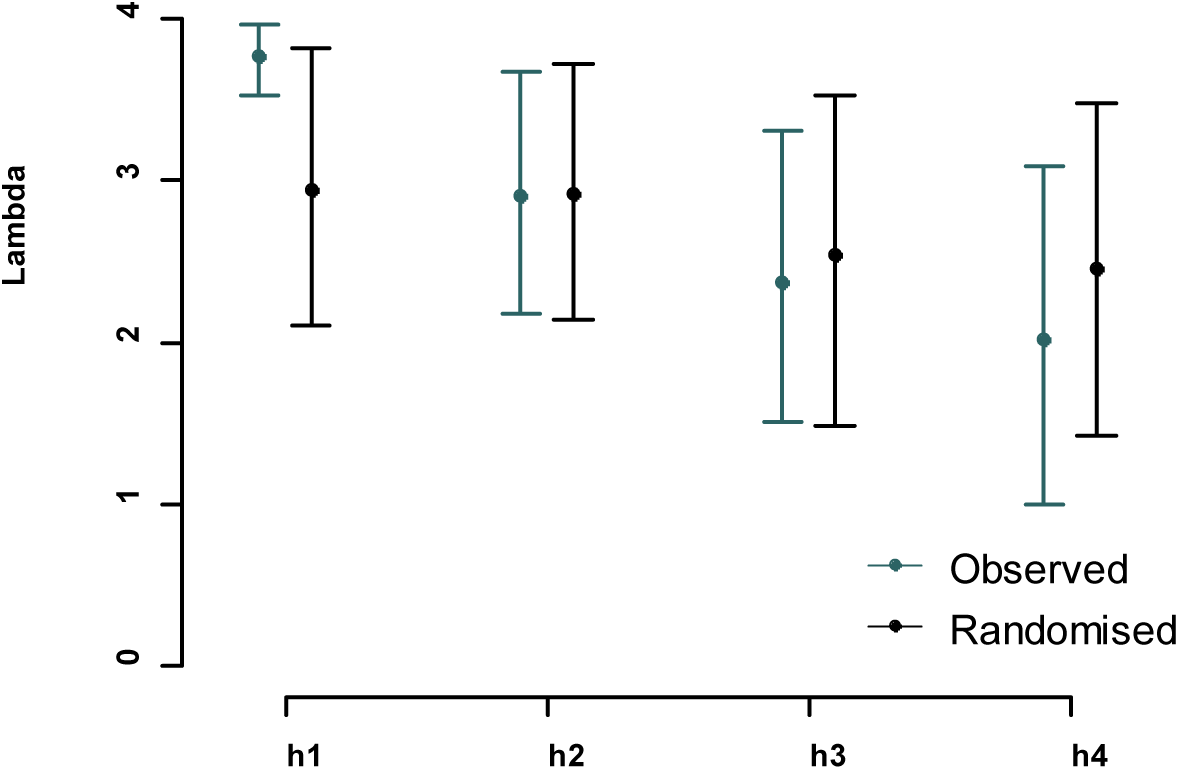
Eigenvalues (lambda) for each eigenvector of the shared subspace **H**, (where h > min k). Higher values of lambda indicate greater similarity of population subspaces.

The second and third eigenvalues of **R** are of similar magnitude, and when projected through the G-matrices, reveal similar patterns. The second axis of **R** captures trait combinations with opposing and larger values for anther-stigma distance and flowering time (Table 2); we observed that Hoffman had more genetic variation for this combination of traits than Pennsylvania. Collectively, these data indicate that the differences in the G-matrices detected by random skewers were primarily driven by the Hoffman population, and it having more variation in the linear combinations of traits measured (*RSe1*: corolla width, flowering time and seed mass in concert; *RSe2*: flowering time and anther stigma distance in opposition).

### Krzanowski’s subspace analysis

To construct **H**, the shared subspace, we used the number of eigenvectors which explained at least 90% of the variation in each population. The first four eigenvectors of each population satisfied this condition. Lambda, which ranges from 1 to 4, approaches 4 in the first eigenvector of **H** (Table 4, Figure 4). A lambda value nearing 4 means the populations all contain the variation sufficient to recreate the shared trait space. Thus, the populations contain considerable shared trait space in the first dimension of **H**. The second through fourth eigenvalues are lower but similar to the randomized comparison (Figure 4). Overall, there is no evidence of divergence among populations in the shared subspace.

**Figure 4:**
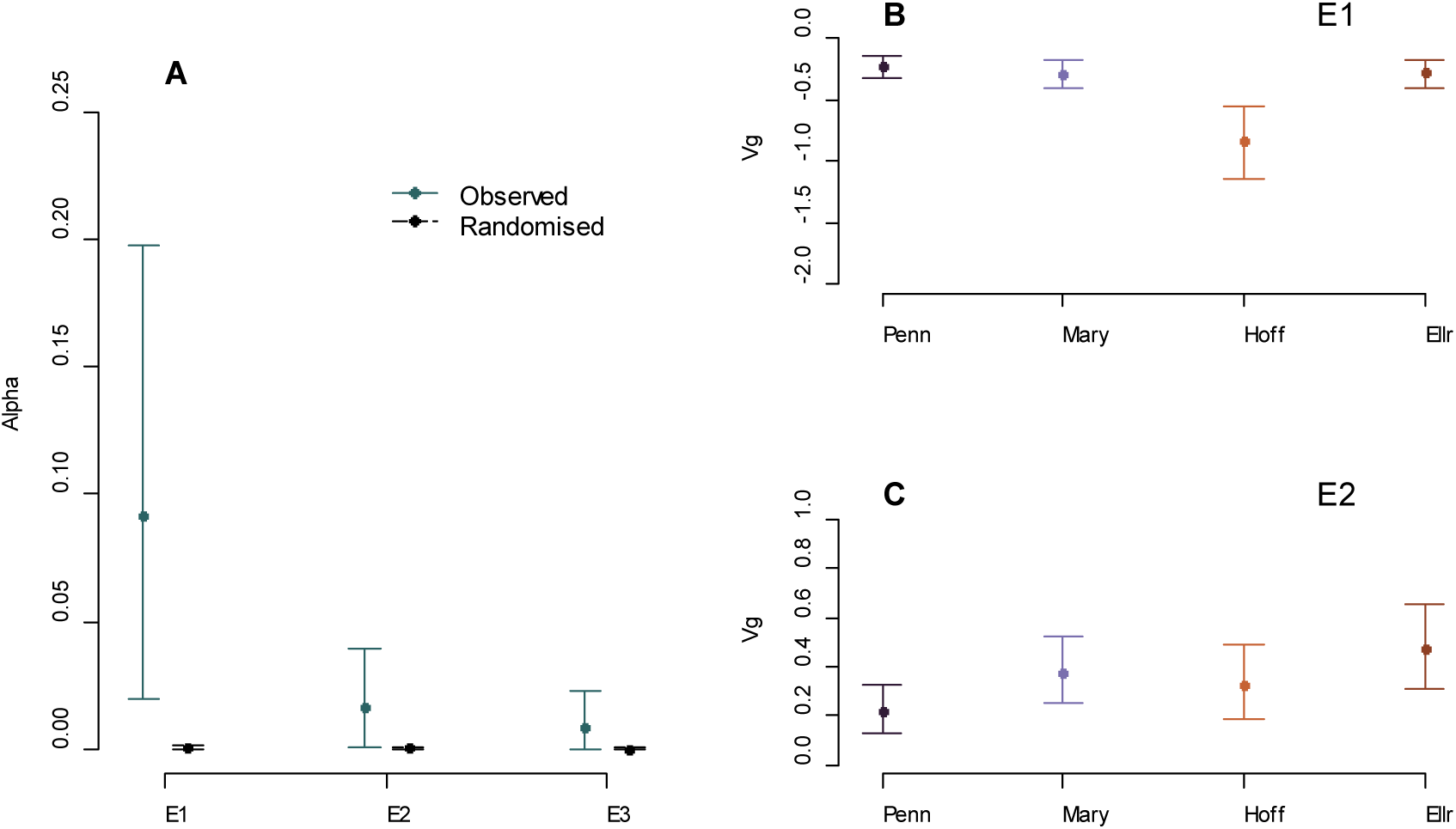
A) Eigenvalues (α) for each non-zero eigentensor from the observed and randomized G- matrices. Note that the upper and lower bounds of the randomized data, reflecting only sampling variation, are small and hard to distinguish on the figure. B) Coordinates of each population in the first non-zero eigentensor of **S**, E1. C) Coordinates of each population in the second non-zero eigentensor of **S,** E2.

**Table 4:** Eigenvalues (lambda) for each eigenvector of the shared subspace **H** with the 95% HPD intervals. A value of 4 would indicate that all the populations have the variation in their subspaces required to recreate the respective eigenvalue of **H**, lower values indicate that populations do not and thus lack similarity.

| Eigenvector | Lambda | Lower HPD | Higher HPD |
| --- | --- | --- | --- |
| **h1** | 3.776 | 3.528 | 3.964 |
| **h2** | 2.909 | 2.184 | 3.679 |
| **h3** | 2.378 | 1.515 | 3.305 |
| **h4** | 2.021 | 0.994 | 3.089 |

To evaluate any population level divergence from the shared subspace we calculated the angle between the axes of **H** and the subspaces of the populations (**A**_t_**A**_t_^T^), which capture at least 90% of the genetic variation in each population. Larger angles indicate that populations differ in some manner along that axis of shared subspace. The first four eigenvectors of Pennsylvania, Hoffman, and Ellerbe, and Maryland were used to generate the shared subspace. Overall, the angles between **H** and the population subspaces are similar and small with increasing uncertainty for higher eigenvectors (Figure S3), meaning differences between the populations and the shared trait subspace are minimal.

### Covariance tensor analysis

We calculated the fourth order covariance tensor for the population G-matrices to assess areas of divergence between the populations. We used the randomized G-matrices, which represent sampling variation within populations, as a null contrast. For observed G-matrices, we estimated three non-zero eigentensors for tensor **S**. Of these three eigentensors, the first two have eigenvalues significantly different from a null expectation based on sampling variation creating each G-matrix independently (Table S6, Figure 4).

To determine the contributions of each population to the eigentensor, we estimated the coordinates of the populations within each eigentensor. The greater the absolute value of the coordinate the greater the contribution a population has. In the first eigentensor (E1, which explains 78.4% of the variation across all non-zero eigentensors), Hoffman again stands out from the other populations and has the greatest contribution. Thus, the first eigentensor, which describes the greatest amount of variation among the population G-matrices, is largely driven by differences between Hoffman, NC and all the other populations. In contrast, Ellerbe, NC has the largest coordinate value in the second eigentensor (E2), which explains 14.4% of the variation among G-matrices.

Eigenanalysis of the eigentensors reveals that the first eigenvector of E1, (*e11*) which explains 45.6% of overall variation (58.11% of variation within E1), is most heavily weighted by seed mass and corolla width (similar to *RSe1* of the R-matrix, as analyzed by Random Skewers), with all trait loadings being of the same sign (Table 5). The first eigenvector of E2 *(e21)*, which explains 6.88% of overall variation among populations (47.17% of variation in E2), describes a dimension most heavily weighted by the seed mass and growth rate of opposing signs (Table 5). Thus these two eigentensors describe population differences in the relationship between seed mass and other traits, especially with respect to the Southern populations.

**Table 5:** Eigenvectors of each non-zero eigentensor of observed **S**. The variance explained by each non- zero eigentensor of **S** is given, along with the eigenvalues of each eigentensor in the direction of each eigenvector. These values describe the contribution of the independent trait combinations described by the eigentensors. The proportion of variance is the proportion of variance of the eigentensors explained by each eigenvector. The trait loadings for eigenvalues of each of the three eigentensor are given, along with the eigenvalue of the eigentensors.

| Eigenvector | $\sigma$ of $E_x$ | Eigenvalue | Prop. of $\sigma$ | Seed mass | Growth | Flow time | Corolla | AS dist. |
| --- | --- | --- | --- | --- | --- | --- | --- | --- |
| e1.1 | 0.784 | -0.914 | 0.581 | -0.618 | -0.134 | -0.372 | -0.607 | -0.306 |
| e1.2 |  | -0.301 | 0.191 | -0.255 | -0.530 | 0.648 | 0.201 | -0.440 |
| e1.3 |  | -0.260 | 0.166 | 0.536 | 0.327 | 0.113 | -0.338 | -0.692 |
| e1.4 |  | 0.072 | 0.046 | 0.513 | -0.770 | -0.306 | -0.207 | 0.085 |
| e1.5 |  | -0.026 | 0.016 | 0.056 | 0.037 | 0.579 | -0.659 | 0.476 |
| e2.1 | 0.144 | 0.840 | 0.471 | 0.677 | -0.642 | -0.309 | -0.105 | 0.150 |
| e2.2 |  | 0.395 | 0.221 | 0.646 | 0.607 | 0.235 | -0.385 | -0.104 |
| e2.3 |  | -0.303 | 0.170 | 0.088 | -0.292 | 0.892 | 0.154 | 0.295 |
| e2.4 |  | 0.216 | 0.121 | -0.220 | 0.104 | -0.129 | -0.503 | 0.819 |
| e2.5 |  | -0.028 | 0.016 | 0.261 | 0.351 | -0.191 | 0.751 | 0.457 |
| e3.1 | 0.071 | 0.675 | 0.355 | 0.948 | -0.092 | 0.001 | 0.288 | -0.102 |
| e3.2 |  | -0.590 | 0.310 | 0.057 | -0.687 | -0.514 | -0.473 | -0.193 |
| e3.3 |  | -0.334 | 0.176 | 0.209 | 0.346 | -0.434 | -0.310 | 0.744 |
| e3.4 |  | -0.291 | 0.153 | -0.153 | -0.597 | 0.083 | 0.520 | 0.585 |
| e3.5 |  | 0.011 | 0.006 | -0.179 | 0.209 | -0.736 | 0.572 | -0.238 |

We next projected these eigenvectors through the population G-matrices, as we did for the random skewers, to assess how much variation each population has along these axes. Not unexpectedly, Hoffman, NC has significantly greater genetic variation along the first and second eigenvector of E1 (Figure 5). Ellerbe has greater variation than the other population along the first eigenvector of E2, although the HPD intervals overlap all other populations. The second eigenvector of E2 similarly has overlapping 95% HPD intervals for all populations but the Southern populations have slightly more genetic variation in this direction than the Northern populations.

**Figure 5:**
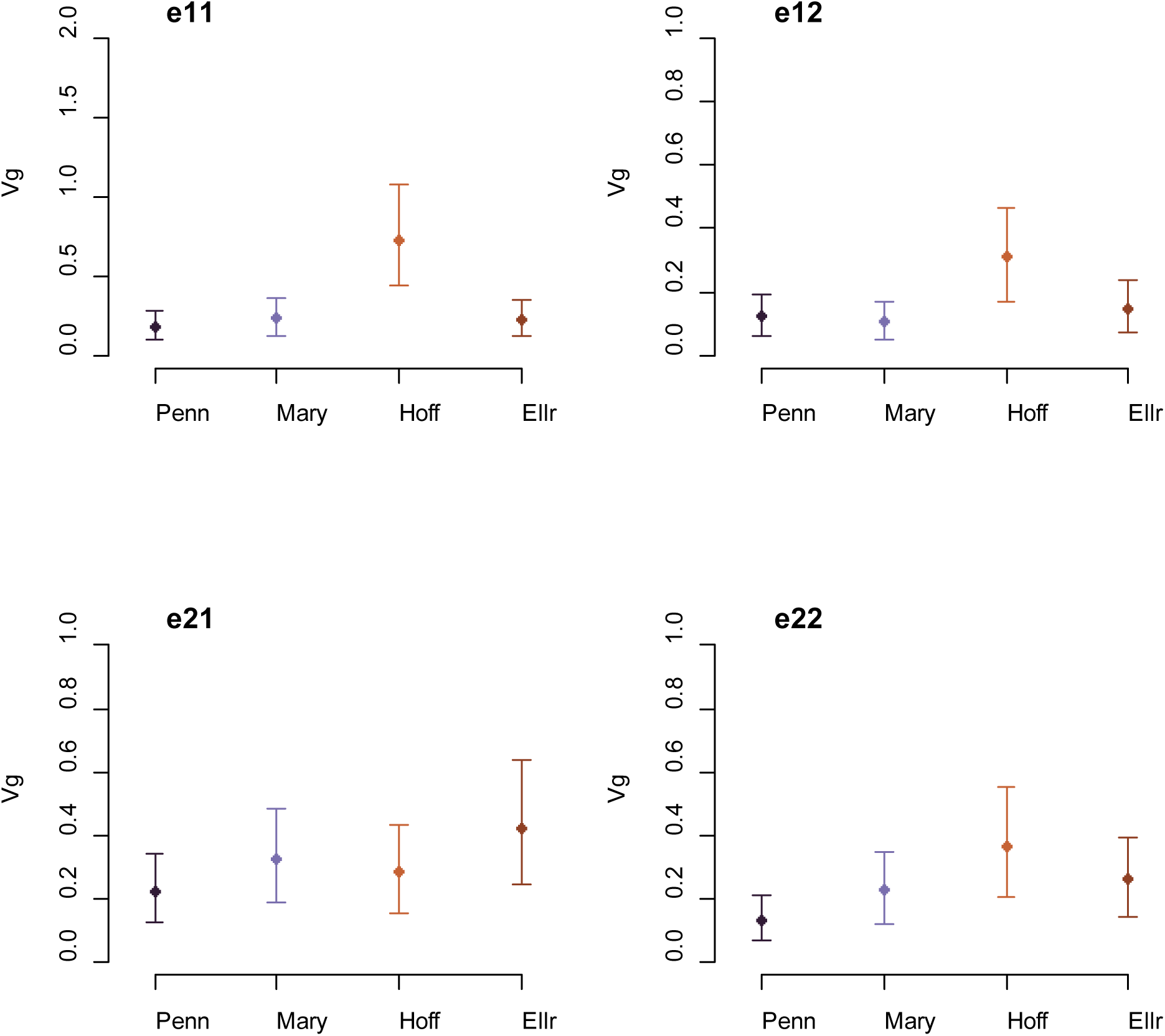
Genetic variation in the direction of eigenvectors of the fourth-order covariance tensor **S**. *e11* is the first eigenvector of the first eigentensor, *e12* is the second eigenvector of the first eigentensor, *e21* is the first eigenvector of the second eigentensor, *e22* is the second eigenvector of the second eigentensor. The vectors were projected through each of the G-matrices to determine the amount of genetic variation that exists in that dimension.

### Similarity between skewers and tensors

Given the similarities between the first eigenvector of E1 of **S** and the results from the random skewers R-matrix projection we calculated the correlation between the two vectors and found that they are almost perfectly correlated (corr = 0.97). This congruence between two vectors of differentiation calculated with entirely different methods provides additional support for this axis describing considerable differences among populations.

## Discussion

Understanding patterns of genetic variation within populations, and how they relate to phenotypic differentiation among them, is a central goal in evolutionary biology. Our analyses of genetic (co)variance matrices and divergence of four natural populations of *Ipomoea hederacea* point to three main results. First, despite the variety of evolutionary forces that could produce differences in **G** between Northern and Southern populations—including strong selection, drift, and bottlenecks—we found remarkably similar patterns of **G** and genetic variation. Our results add to a growing body of cases in which differences in **G** are predicted *a priori*, but not detected. Second, contrary to a variety of theoretical predictions, we failed to detect a relationship between the axis of greatest multivariate clinal divergence and the axes of greatest genetic variation within populations. Below, we discuss these results in light of how genetic constraints and reduced dimensionality of **G** influence phenotypic divergence, and divergence in **G** these populations.

### Evolutionary potential and divergence of G

Our study of **G** within these four populations allows both an evaluation of the evolutionary potential within each population, and the divergence (or lack thereof) of **G** itself between range-edge populations and central populations. Using a variety of approaches, including estimating trace of **G**, and three methods for comparing **G** among populations, we found a consistent pattern of results. These populations all appear to be of reduced rank, have similar amounts of total genetic variance, and there do not appear to be appreciable differences between range-edge populations and core populations in the structure, size, or orientation of **G**.

We first assessed the total amount of genetic variation among populations and found that there was a trend along latitude for decreased genetic variation in the Northern populations, although the 95% HPD intervals of one of the Southern populations overlapped, making this trend slight. Hoffman, NC, one of the Southern populations, stood out among others as possessing significantly greater variation than all other populations. Similarly, one of the Northern populations, Pennsylvania, had the lowest trace, and if it had been estimated as slightly smaller (15% reduction), its trace would not have overlapped with the Southern populations.

However, while we detected one Southern population with increased genetic variation, and possibly one Northern population with reduced variation, the remaining two populations have robustly similar amounts of quantitative genetic variation. In like fashion, trait heritabilities tended to be higher in Southern populations (Table S1), albeit again with overlapping HPD intervals. Thus, it appears that the Hoffman population is distinct from the rest, rather than Southern populations overall strongly differing from Northern populations.

Two of the traits, corolla width and anther-stigma distance, had very small variances indistinguishable from sampling variance in three of the populations (Hoffman being the exception). The lack of variation in these traits in **G** make observing differences among populations impossible with respect to corolla width and anther-stigma distance. In Cline_max_ both corolla width and anther-stigma distance showed significant among-population divergence of the means, which suggests that the paucity of within-population variation for these traits is likely due to selection. We retained these traits in our G-matrices to allow for comparison with the axis of greatest multivariate divergence, Cline_max_ (from Stock et al. 2014).

Our results are suggestive of the overall stability of the G-matrices across populations, and add to a handful of cases in which **G** remains similar despite adaptation to gradients, differences in selective regimes, and altered patterns of gene flow. Given the *a priori* expectation that drift and population bottlenecks should introduce stochastic changes in **G** (Lande 1979, Shaw et al. 1995, Roff 2000, Phillips et al. 2001, Jones et al. 2004, Doroszuk et al. 2008) and that populations of *Ipomoea hederacea* likely have low effective populations sizes and may frequently experience bottlenecking (Campitelli & Stinchcombe 2014) and thus are expected to be susceptible to drift, the degree of stability we found is surprising. In this regard, our results are similar to McGoey and Stinchcombe (2021), who found little difference in G-matrices between introduced and native populations of ragweed (*Ambrosia artemisiifolia*) despite expectations to the contrary. More generally, our results add to other findings of **G** being relatively stable across geography (Arnold et al. 2008; Delahaie et al. 2017), even in cases in which theoretical guidance suggests a strong expectation of divergence in G. In Scandinavian populations of *Arabidopsis thaliana*, Puentes et al. (2016) expected to find divergence in **G** due to differences in selective environments and limited gene flow. While the populations had univariate differences the G- matrices of all populations demonstrated overall similarity in shape and orientation. Similarly, Hangartner et al. (2019) estimated **G** in locally adapted populations along a broad climatic gradient with disparate selective regimes and found that **G** had minimal changes across eastern Australian populations of *Drosophila melanogaster*. In island and mainland populations of the blue tit (*Cyanistes caeruleus*) where two distinct, heterogenous landscapes produce divergent selection and gene flow between populations is expected to be low to effectively absent (in the case of island-mainland) Delahaie et al. (2017) found weak overall differentiation in **G**. Although **G** can evolve under certain conditions, and empirical studies have in some cases detected remarkable differences, ours is one of the growing number of studies that have failed to detect changes in **G** despite strong expectations.

Both the covariance tensor and random skewers analysis suggest that the Hoffman, NC population differed from the others (this population also had the most genetic variation overall and in the direction of clinal divergence). The covariance tensor and skewers approaches are complementary, in that the tensor reflects patterns of variation and covariation among **G**, while the skewers analysis is insensitive to differences in the total amount of genetic variance and magnitude of the response (as it uses the correlation between response vectors, but not their magnitude; Hansen and Houle 2008). These differences are not simply due to Hoffman’s greater overall genetic variation. The first eigenvector of the first eigentensor (*e11*), the first eigenvector of the random skewers analysis (*RSe1*), and g_max_ within the Hoffman population were all highly correlated, with heavy loading from flowering time, corolla width, and seed mass. The Hoffman, NC population site did not differ substantially from the others: all were collected from roadsides adjacent to agricultural fields and were reasonably large to allow for sampling from 70-100 individuals. Hoffman was, however, the only population where we observed both leaf shape genotypes (leaf shape is a simple Mendelian polymorphism in this species; Bright-Emlen 1998). Some speculative possibilities are that it was founded by heterozygous individuals, multiple founders of opposite leaf shape genotypes, or is subject to balancing selection on the leaf shape locus (as has been shown by Bright and Rausher 2008), and thus has more variation for any of these reasons.

Collectively, our diverse set of analyses points to a single population being qualitatively different from the others, with divergence driven by a handful of traits in that population, rather than range-edge populations having any inherent differences in the structure of **G**. At face value, these findings might suggest that range-edge populations of *Ipomoea hederacea* are not small enough to experience substantial genetic drift that could reduce genetic variance compared to central populations; alternatively, it may be that all populations experience frequent extinction and colonization dynamics, across the range, and that this does not have differential effects on genetic variance. We suspect the latter possibility is more likely, especially when taken together with the population genetic work of Campitelli and Stinchcombe (2014), in which they suggested that metapopulation dynamics with high extinction and dispersal rates may explain the patterns of neutral genetic diversity.

### Clinal divergence and G

In general, phenotypic divergence is expected to be aligned with **G** under three overall conditions. First, under genetic drift, we expect that phenotypic divergence in a suite of traits should be related to the overall amount of standing quantitative genetic variance in those traits (Lande 1979, Phillips et al. 2001). Second, when G-matrices are ill-conditioned, or of reduced rank, we expect short-term evolutionary responses to be biased towards directions of multivariate trait space with genetic variance (Chenoweth et al. 2010). In other words, once a G-matrix differs from spherical, evolutionary responses will be dominated by the directions containing the most variance. In the third scenario, when β = g_max_, divergence, Δz, proceeds along g_max_, and we expect the maximal evolutionary response (Gaydos et al. 2013). Hangartner et al. (2019) found that G-matrices of range-edge populations of *Drosophila melanogaster* were in fact aligned with clinal divergence and that the trait covariances improved the adaptive potential of peripheral populations. Work on the cuticular hydrocarbons of *Drosophila serrata* population by Hine and colleagues (Hine et al 2009, Chenoweth et al. 2008) found patterns in mean trait divergence, reduction in associated variation in g_max_, and divergence among populations along g_max_. Similarly, Aguirre et al (2014) found that those populations share considerable trait space, although there were axes of divergence among the populations. In contrast, phenotypic divergence does not have to be aligned with **G** if selection has been strong and persistently favoring alternative combinations of traits, or consistent selection over long evolutionary timescales. For example, McGuigan et al. (2005), found that hydrodynamic adaptation in rainbow fish was only weakly associated with g_max_, and was primarily aligned with the trailing eigenvectors of **G**; they interpreted this as a result of long-term selection towards a new phenotypic optimum.

With this context in mind, many of our results suggest that we should have observed a relationship between clinal divergence and **G**: the traits are genetically variable, both individually (Table S1) and as a linear combination (Table 2). It also seems unlikely that the phenotypic divergence detected is a result of long-term selection, similar to McGuigan et al. (2005), as *I. hederacea* currently inhabits (and was collected from) ephemeral, disturbance-prone habitats such as roadsides, agricultural fields, and cleared areas. Consequently, many of the conditions that can produce a relationship between **G** and divergence appear to be met.

Two potential explanations for why we do not see such a relationship between **G** and divergence occur to us. The first is that previous work on *Ipomoea hederacea* (Simonsen and Stinchcombe 2010; Campitelli and Stinchcombe 2013; Stock et al. 2014) and this study inadvertently omitted an ecologically important, correlated trait. For this scenario to explain the lack of relationship between divergence and **G**, there would have to be an omitted trait that is both highly variable within populations, and highly diverged between populations, such that its inclusion would predominate both g_max_ and any estimate of a divergence vector. The problem of missing traits is inherent to studies of phenotypic evolution; while the traits we measured capture aspects of size, growth, phenology, and floral morphology, it is possible that other traits related to ecophysiology, seed bank dynamics, or other aspects of the life cycle are more important.

Second, it may be that natural selection is acting in a direction that is nearly orthogonal to g_max_, thus leading to the divergence that is at a substantial angle from g_max_. If this were the case, it would be possible for the G-matrices we estimated to produce evolutionary responses that could lead to the observed divergence. One clue in support of this hypothesis is that flowering time appears to load relatively weakly on g_max_ in the Pennsylvania population, but in the past has been detected to be under very strong directional selection in this species (Simonsen and Stinchcombe 2010; Campitelli and Stinchcombe 2013). Similarly, Campitelli and Stinchcombe (2013) found selection favoring decreased flowering time and increased early and mid-season growth rates (i.e., β elements of opposite signs), while we found that growth rate and flowering time load in the same direction (i.e., are positively correlated) of g_max_ in three of four populations. As such, there are past observations of selection acting strongly on traits with relatively little genetic variance, or in opposite directions on positively correlated traits, both of which could lead to β being orthogonal to g_max_. Indeed, Simonsen and Stinchcombe (2010) found strong natural selection almost orthogonal to g_max_ in a field study of size traits and flowering time, due to size traits being highly variable and under weak selection, and flowering time being less variable but under very strong selection. The divergence observed may thus be the product of selection, but without requiring recourse to long-term selection towards a single optimum. While there are field studies of natural selection on some of the traits we studied, an overall study of all of them is necessary to distinguish between the missing trait explanation and **β** being orthogonal to g_max_.

### Conclusions and Future Directions

Determining the relationship between **G** and phenotypic divergence, or divergence of G- matrices themselves, is a challenging empirical and statistical endeavor. Practical constraints make constructing well estimated G-matrices for dozens of populations nearly impossible (Arnold et al. 2008) is certainly the case for *I. hederacea*. The most tractable and testable hypothesis to emerge from our work is that natural selection on these five traits is poorly aligned with g_max_. Future field studies will be required to further elucidate the action of natural selection on this species, and the ecological mechanisms behind selection for this suite of traits and others.

## References

1. Agrawal, A. F., and J. R. Stinchcombe. 2009. How much do genetic covariances alter the rate of adaptation? Proc. Biol. Sci. 276:1183–1191.

2. Aguirre, J. D., E. Hine, K. McGuigan, and M. W. Blows. 2014. Comparing **G**: multivariate analysis of genetic variation in multiple populations. Heredity 112:21–29.

3. Antonovics, J. 1976. The Nature of Limits to Natural Selection. Ann. Mo. Bot. Gard. 63:224–247.

4. Arnold, S. J., R. Bürger, P. A. Hohenlohe, B. C. Ajie, and A. G. Jones. 2008. Understanding the evolution and stability of the G-matrix. Evolution 62:2451–2461.

5. Barton, N. H. 2001. Adaptation at the edge of a species’ range. Pp. 365–392 in Integrating Ecology and Evolution in a Spatial Context, Silvertown J and Antonovics, ed.. Cambridge University Press.

6. Barton, N. H., and M. Turelli. 1987. Adaptive landscapes, genetic distance and the evolution of quantitative characters. Genet. Res. 49:157–173.

7. Basser, P. J., and S. Pajevic. 2007. Spectral decomposition of a 4th-order covariance tensor: Applications to diffusion tensor MRI. Signal Processing 87:220–236.

8. Björklund, M., A. Husby, and L. Gustafsson. 2013. Rapid and unpredictable changes of the G-matrix in a natural bird population over 25 years. J. Evol. Biol. 26:1–13.

9. Bright-Emlen, K. L. 1998. Geographic variation and natural selection on a leaf shape polymorphism in the ivyleaf morning glory (Ipomoea hederacea). Duke University, Ann Arbor, United States.

10. Bright, K. L., and M. D. Rausher. 2008. Natural selection on a leaf-shape polymorphism in the ivyleaf morning glory (*Ipomoea hederacea*). Evolution 62:1978–1990.

11. Brown, J. H., D. W. Mehlman, and G. C. Stevens. 1995. Spatial variation in abundance. Ecology 76:2028–2043.

12. Campitelli, B. E., and J. R. Stinchcombe. 2013. Natural selection maintains a single-locus leaf shape cline in Ivyleaf morning glory, *Ipomoea hederacea*. Mol. Ecol. 22:552–564.

13. Campitelli, B. E., and J. R. Stinchcombe. 2014. Population dynamics and evolutionary history of the weedy vine *Ipomoea hederacea* in North America. G3 4:1407–1416.

14. Chantepie, S., and L.-M. Chevin. 2020. How does the strength of selection influence genetic correlations? Evol. Lett. 4:468–478.

15. Chenoweth, S. F., H. D. Rundle, and M. W. Blows. 2008. Genetic constraints and the evolution of display trait sexual dimorphism by natural and sexual selection. Am. Nat. 171:22– 34.

16. Chenoweth, S. F., H. D. Rundle, and M. W. Blows. 2010. The contribution of selection and genetic constraints to phenotypic divergence. Am. Nat. 175:186–196.

17. Cheverud, J. M., and G. Marroig. 2007. Comparing covariance matrices: Random skewers method compared to the common principal components model. Genetics and Molecular Biology 30:461–469.

18. Conner, J. K., K. Karoly, C. Stewart, V. A. Koelling, H. F. Sahli, and F. H. Shaw. 2011. Rapid independent trait evolution despite a strong pleiotropic genetic correlation. Am. Nat. 178:429–441.

19. Costa E, Silva J., B. M. Potts, and P. A. Harrison. 2020. Population divergence along a genetic line of least resistance in the tree species *Eucalyptus globulus*. Genes 11:1095.

20. Delahaie, B., A. Charmantier, S. Chantepie, D. Garant, M. Porlier, and C. Teplitsky. 2017. Conserved G-matrices of morphological and life-history traits among continental and island blue tit populations. Heredity 119:76–87.

21. Doroszuk, A., M. W. Wojewodzic, G. Gort, and J. E. Kammenga. 2008. Rapid divergence of genetic variance-covariance matrix within a natural population. Am. Nat. 171:291–304.

22. Endler, J. A. 1977. Geographic Variation, Speciation and Clines. Princeton University Press.

23. Ennos, R. A. 1981. Quantitative studies of the mating system in two sympatric species of *Ipomoea* (Convolvulaceae). Genetica 57:93–98.

24. Falconer, D. S., and T. F. C. Mackay. 1996. Introduction to quantitative genetics. Fourth. Essex, UK: Longman Group.

25. Garcia-Ramos, G., and M. Kirkpatrick. 1997. Genetic models of adaptation and gene flow in peripheral populations.

26. Gaydos, T. L., N. E. Heckman, M. Kirkpatrick, J. R. Stinchcombe, J. Schmitt, J. Kingsolver, and J. S. Marron. 2013. Visualizing genetic constraints. Ann. Appl. Stat. 7:860–882.

27. Guillaume, F., and M. C. Whitlock. 2007. Effects of migration on the genetic covariance matrix. Evolution 61:2398–2409.

28. Hadfield, J. D. 2010. MCMC Methods for Multi-Response Generalized Linear Mixed Models: The MCMCglmm R Package. J. Stat. Softw. 33:1–22.

29. Hangartner, S., Lasne, C., Sgrò C.M., Connallon T., and K. Monro. 2019. Genetic covariances promote climatic adaptation in Australian *Drosophila*. Evolution 74: 326–337.

30. Hansen, T. F., and D. Houle. 2008. Measuring and comparing evolvability and constraint in multivariate characters. J. Evol.

31. Hine, E., S. F. Chenoweth, H. D. Rundle, and M. W. Blows. 2009. Characterizing the evolution of genetic variance using genetic covariance tensors. Philos. Trans. R. Soc. Lond. B Biol. Sci. 364:1567–1578.

32. Hull-Sanders, H. M., M. D. Eubanks, and D. E. Carr. 2005. Inbreeding depression and selfing rate of *Ipomoea hederacea* var. *integriuscula* (Convolvulaceae). Am. J. Bot. 92:1871– 1877.

33. Johnson, T., and N. Barton. 2005. Theoretical models of selection and mutation on quantitative traits. Philos. Trans. R. Soc. Lond. B Biol. Sci. 360:1411–1425.

34. Jones, A. G., S. J. Arnold, and R. Bürger. 2004. Evolution and stability of the G-matrix on a landscape with a moving optimum. Evolution 58:1639–1654.

35. Kingsolver, J. G., N. Heckman, J. Zhang, P. A. Carter, J. L. Knies, J. R. Stinchcombe, and K. Meyer. 2015. Genetic variation, simplicity, and evolutionary constraints for function- valued traits. Am. Nat. 185:E166–81.

36. Kirkpatrick, M. 2009. Patterns of quantitative genetic variation in multiple dimensions. Genetica 136:271–284.

37. Kirkpatrick, M., and N. H. Barton. 1997. Evolution of a species’ range. Am. Nat. 150:1–23.

38. Klingaman, T. E., and L. R. Oliver. 1996. Existence of ecotypes among populations of entireleaf morningglory (*Ipomoea hederacea* var. *integriuscula*). Weed Sci. 44:540–544.

39. Krzanowski, W. J. 1979. Between-groups comparison of principal components. J. Am. Stat. Assoc. 74:703–707.

40. Lande, R. 1992. Neutral theory of quantitative genetic variance in an island model with local extinction and colonization. Evolution 46:381–389.

41. Lande, R. 1979. Quantitative genetic analysis of multivariate evolution, applied to brain: body size allometry. Evolution 33:402–416.

42. Lande, R., and S. J. Arnold. 1983. The measurement of selection on correlated characters. Evolution 37:1210–1226.

43. Lin, C. Y., and F. R. Allaire. 1977. Heritability of a linear combination of traits. Theor. Appl. Genet. 51:1–3.

44. McGoey, B. V., and J. R. Stinchcombe. 2021. Introduced populations of ragweed show as much evolutionary potential as native populations. Evol. Appl. 14:1436–1449.

45. McGuigan, K., S. F. Chenoweth, and M. W. Blows. 2005. Phenotypic divergence along lines of genetic variance. Am. Nat. 165:32–43.

46. Morrissey, M. B., S. Hangartner, and K. Monro. 2019. A note on simulating null distributions for G-matrix comparisons. Evolution 73:2512–2517.

47. Peterson, R. A., and J. E. Cavanaugh. 2020. Ordered quantile normalization: a semiparametric transformation built for the cross-validation era. J. Appl. Stat. 47:2312–2327.

48. Phillips, P. C., M. C. Whitlock, and K. Fowler. 2001. Inbreeding changes the shape of the genetic covariance matrix in *Drosophila melanogaste*r. Genetics 158:1137–1145.

49. Polechová, J., and N. H. Barton. 2015. Limits to adaptation along environmental gradients. PNAS 112:6401–6406.

50. Polechová, J. 2018. Is the sky the limit? On the expansion threshold of a species’ range. PLoS Biol. 16:e2005372.

51. Puentes, A., G. Granath, and J. Ågren. 2016. Similarity in G-matrix structure among natural populations of *Arabidopsis lyrata*. Evolution 70:2370–2386.

52. Roff, D. 2000. The evolution of the G-matrix: selection or drift? Heredity 84:135–142.

53. Roff, D. and T.A. Mousseau. 1999. Does natural selection alter genetic architecture? An evaluation of quantitative genetic variation among populations of Allenomobius socius and A. fasciatus. J. Evol. Biol. 12:361–369.

54. Roughgarden, J. 1979. Theory of population genetics and evolutionary ecology: an introduction. Macmillan, New York.

55. Royauté, R., A. Hedrick, and N. A. Dochtermann. 2020. Behavioural syndromes shape evolutionary trajectories via conserved genetic architecture. Proc. Biol. Sci. 287:20200183.

56. Sexton, J. P., P. J. McIntyre, A. L. Angert, and K. J. Rice. 2009. Evolution and ecology of species range limits. Annu. Rev. Ecol. Evol. Syst. 40:415–436.

57. Simonsen, A. K., and J. R. Stinchcombe. 2010. Quantifying evolutionary genetic constraints in the ivyleaf morning glory, *ipomoea hederacea*. Int. J. Plant Sci. 171:972–986.

58. Slatkin, M. 1978. Spatial patterns in the distributions of polygenic characters. J. Theor. Biol. 70:213–228.

59. Stock, A. J., B. E. Campitelli, and J. R. Stinchcombe. 2014. Quantitative genetic variance and multivariate clines in the Ivyleaf morning glory, *Ipomoea hederacea*. Philos. Trans. R. Soc. Lond. B Biol. Sci. 369:20130259.

60. Shaw, F. H., R. G. Shaw, G. S. Wilkinson, and M. Turelli. 1995. Changes in genetic variances and covariances: **G** whiz! Evolution 49:1260–1267.

61. Sztepanacz, J. L., and M. W. Blows. 2017. Accounting for sampling error in genetic eigenvalues using random matrix theory. Genetics 206:1271–1284.

62. Teplitsky, C., M. R. Robinson, and J. Merilä. 2014. Evolutionary potential and constraints in wild populations. Quantitative genetics in the wild 190–208. Oxford University Press Oxford, UK.

63. Turelli, M. 1988. Phenotypic evolution, constant covariances, and the maintenance of additive variance. Evolution 42:1342–1347.

64. Walsh, B., and M. W. Blows. 2009. Abundant genetic variation + strong selection = multivariate genetic constraints: a geometric view of adaptation. Annu. Rev. Ecol. Evol. Syst. 40:41–59.

65. Uesugi, A., T. Connallon, A. Kessler, and K. Monro. 2017. Relaxation of herbivore-mediated selection drives the evolution of genetic covariances between plant competitive and defense traits. Evolution 71:1700–1709.

66. Walter, G. M., J. David Aguirre, M. W. Blows, and D. Ortiz-Barrientos. 2018. Evolution of genetic variance during adaptive radiation. Am. Nat. 191:E108–E128.

67. Whitlock, M. C., and D. E. McCauley. 1990. Some population genetic consequences of colony formation and extinction: genetic correlations within founding groups. Evolution 44:1717–1724.

68. Wood, C. W., and E. D. Brodie 3rd. 2015. Environmental effects on the structure of the G-matrix. Evolution 69:2927–2940.

