## Supplementary materials for "G-matrix stability in clinally diverging populations of an annual weed"

Univariate analysis

In addition to the measured traits we calculated a composite trait, the index value, using mean-standardized trait values weighted by the coefficients of the axis of greatest multivariate clinal divergence (Cline_max_) from Stock et al. (2014):

${Index}_{i}=\sum_{j}^{5} {Trait}_{ij}* {Coeff}_{j}$

Where the index value for the *i* ^th^ individual is equal to the sum of each of the *i* ^th^ individual’s *j* ^th^ trait value multiplied by the *j* ^th^ trait coefficient of Cline_max_. More positive values indicate more northern multivariate trait values. Here, we use the mean-standardized Cline_max_ as reported in Stock et al. (2014). We note that in all multivariate analyses we have use a standard deviation-standardized Cline_max_.

The index value follows the same general trend as in individual traits, with lower heritability estimates in northern populations (Pennsylvania: 0.046, Maryland: 0.121, Hoffman, NC: 0.163, Ellerby, NC: 0.187) but with overlapping HPD intervals (Table S1). The northern populations also tended to have more positive mean estimated values, suggesting the populations in this analysis are consistent with those used in Stock et al. (2014). One of the northern populations, Maryland, had an estimated mean trait value which was significantly greater than all other populations (Table S1).

**Table S1**: Results from univariate Bayesian models. Each trait was modelled with all populations as fixed effects, and with the environmental block (maternal greenhouse block in the case of seed mass) and family as random effects. All Highest Posterior Density estimates are 95% intervals.

| **Trait** | **Population** | **Mean** | **SE** | **Dam** σ | **Dam ΔDIC** | **H**² | **HPD Low** | **HPD High** | **No. Dams** | **No. Plants** |
| --- | --- | --- | --- | --- | --- | --- | --- | --- | --- | --- |
| SM.s | PENN | 0.63 | 0.03 | 0.247 | -217.1 | 0.371 | 0.2319 | 0.5314 | 50 | 539 |
| SM.s | MARY | 0.28 | 0.04 | 0.386 | -244.2 | 0.422 | 0.3099 | 0.5415 | 50 | 543 |
| SM.s | HOFF | -0.58 | 0.04 | 0.532 | -359.3 | 0.535 | 0.3529 | 0.6903 | 50 | 530 |
| SM.s | ELLR | -0.35 | 0.04 | 0.387 | -291.7 | 0.499 | 0.3569 | 0.6226 | 49 | 525 |
| GR.s | PENN | 0.25 | 0.04 | 0.071 | -19.9 | 0.081 | 0.0162 | 0.1556 | 50 | 539 |
| GR.s | MARY | -0.01 | 0.05 | 0.143 | -45.9 | 0.116 | 0.0405 | 0.2024 | 50 | 543 |
| GR.s | HOFF | 0.1 | 0.04 | 0.242 | -104.3 | 0.228 | 0.1131 | 0.3498 | 50 | 530 |
| GR.s | ELLR | -0.35 | 0.04 | 0.282 | -124.8 | 0.289 | 0.1786 | 0.3983 | 49 | 525 |
| FT.s | PENN | 0.13 | 0.04 | 0.094 | -148.9 | 0.114 | 0.0369 | 0.2276 | 50 | 539 |
| FT.s | MARY | -1.15 | 0.02 | 0.032 | -56.9 | 0.085 | 0.0270 | 0.1570 | 50 | 543 |
| FT.s | HOFF | 0.19 | 0.03 | 0.239 | -365.8 | 0.464 | 0.3139 | 0.6134 | 50 | 530 |
| FT.s | ELLR | 0.87 | 0.03 | 0.070 | -73.8 | 0.165 | 0.0662 | 0.2623 | 49 | 525 |
| CW.s | PENN | -0.11 | 0.04 | 0.014 | 0.1 | 0.013 | 0.0003 | 0.0440 | 50 | 539 |
| CW.s | MARY | -0.02 | 0.04 | 0.152 | -45.9 | 0.148 | 0.0616 | 0.2342 | 50 | 543 |
| CW.s | HOFF | -0.08 | 0.05 | 0.349 | -131.1 | 0.309 | 0.1903 | 0.4236 | 50 | 530 |
| CW.s | ELLR | 0.21 | 0.04 | 0.116 | -37.9 | 0.124 | 0.0520 | 0.2066 | 49 | 525 |
| ASD.s | PENN | 0.18 | 0.04 | 0.011 | 0.8 | 0.011 | 0.0002 | 0.0363 | 50 | 539 |
| ASD.s | MARY | -0.24 | 0.04 | 0.071 | -23.3 | 0.089 | 0.0213 | 0.1592 | 50 | 543 |
| ASD.s | HOFF | 0.14 | 0.05 | 0.292 | -92.2 | 0.251 | 0.1386 | 0.3623 | 50 | 530 |
| ASD.s | ELLR | -0.08 | 0.04 | 0.104 | -23.9 | 0.097 | 0.0224 | 0.1668 | 49 | 525 |
| Index | PENN | -0.05 | 0.01 | 0.002 | -15.2 | 0.046 | 0.0059 | 0.0977 | 50 | 539 |
| Index | MARY | 0.20 | 0.01 | 0.005 | -70.1 | 0.121 | 0.0519 | 0.1961 | 50 | 543 |
| Index | HOFF | -0.09 | 0.01 | 0.007 | -91.6 | 0.163 | 0.0729 | 0.2550 | 50 | 530 |
| Index | ELLR | -0.06 | 0.01 | 0.008 | -103.1 | 0.187 | 0.0927 | 0.2831 | 49 | 525 |

**Table S2:** Results from univariate REML models. Model structure was the same as the Bayesian models.

| **Trait** | **Population** | **Mean** | **SE** | **F-value** | **Pop p-value** | **Sign. diff** | **Dam** σ | **Dam p-value** | **H**² |
| --- | --- | --- | --- | --- | --- | --- | --- | --- | --- |
| SM.s | PENN | 0.63 | 0.03 | 36.85 | 0 | c | 0.137 | 0 | 0.224 |
| SM.s | MARY | 0.28 | 0.04 | 36.85 | 0 | b | 0.185 | 0 | 0.212 |
| SM.s | HOFF | -0.58 | 0.04 | 36.85 | 0 | a | 0.275 | 0 | 0.3 |
| SM.s | ELLR | -0.35 | 0.04 | 36.85 | 0 | a | 0.194 | 0 | 0.266 |
| GR.s | PENN | 0.25 | 0.04 | 11.5 | 0 | b | 0.06 | 0.013 | 0.073 |
| GR.s | MARY | -0.01 | 0.05 | 11.5 | 0 | b | 0.153 | 0 | 0.138 |
| GR.s | HOFF | 0.1 | 0.04 | 11.5 | 0 | b | 0.169 | 0 | 0.173 |
| GR.s | ELLR | -0.35 | 0.04 | 11.5 | 0 | a | 0.138 | 0 | 0.148 |
| FT.s | PENN | 0.13 | 0.04 | 227.55 | 0 | b | 0.358 | 0 | 0.419 |
| FT.s | MARY | -1.15 | 0.02 | 227.55 | 0 | a | 0.071 | 0 | 0.196 |
| FT.s | HOFF | 0.19 | 0.03 | 227.55 | 0 | b | 0.126 | 0 | 0.28 |
| FT.s | ELLR | 0.87 | 0.03 | 227.55 | 0 | c | 0.001 | 0 | 0.003 |
| CW.s | PENN | -0.11 | 0.04 | 4.84 | 0.003 | a | 0.107 | 0 | 0.112 |
| CW.s | MARY | -0.02 | 0.04 | 4.84 | 0.003 | ab | 0.166 | 0 | 0.166 |
| CW.s | HOFF | -0.08 | 0.05 | 4.84 | 0.003 | ab | 0.305 | 0 | 0.276 |
| CW.s | ELLR | 0.21 | 0.04 | 4.84 | 0.003 | b | 0.091 | 0 | 0.1 |
| ASD.s | PENN | 0.18 | 0.04 | 12.12 | 0 | c | 0.011 | 0.5 | 0.011 |
| ASD.s | MARY | -0.24 | 0.04 | 12.12 | 0 | a | 0.124 | 0 | 0.162 |
| ASD.s | HOFF | 0.14 | 0.05 | 12.12 | 0 | bc | 0.259 | 0 | 0.23 |
| ASD.s | ELLR | -0.08 | 0.04 | 12.12 | 0 | ab | 0.061 | 0.033 | 0.058 |
| Index | PENN | -0.05 | 0.01 | 78.42 | 0 | a | 0.003 | 0 | 0.085 |
| Index | MARY | 0.20 | 0.01 | 78.42 | 0 | b | 0.003 | 0 | 0.085 |
| Index | HOFF | -0.09 | 0.01 | 78.42 | 0 | a | 0.007 | 0 | 0.208 |
| Index | ELLR | -0.06 | 0.01 | 78.42 | 0 | a | 0.009 | 0 | 0.254 |

**Table S3:** **G** matrices estimated for each population along with 95% HPD intervals. Bolded values indicate those which do not overlap the null **G** 95% HPD intervals.

| **Pennsylvania** | **SM** | **95% HPD** | **GR** | **95% HPD2** | **FT** | **95% HPD3** | **CW** | **95% HPD4** | **ASD** | **95% HPD5** |
| --- | --- | --- | --- | --- | --- | --- | --- | --- | --- | --- |
| SM | **0.2747** | **0.147, 0.379** | |  |  |  |  |  |  |  |
| GR | -0.0315 | -0.094, 0.038 | 0.1414 | 0.035, 0.163 | |  |  |  |  |  |
| FT | -0.0068 | -0.066, 0.049 | -0.0195 | -0.064, 0.022 | **0.1092** | **0.055, 0.154** | |  |  |  |
| CW | 0.0170 | -0.039, 0.069 | 0.0227 | -0.012, 0.061 | 0.0100 | -0.026, 0.043 | 0.0900 | 0.012, 0.074 | |  |
| ASD | 0.0081 | -0.044, 0.062 | -0.0103 | -0.047, 0.023 | 0.0067 | -0.021, 0.049 | 0.0041 | -0.022, 0.024 | 0.0925 | 0.009, 0.060 |
| **Maryland** | **SM** | **95% HPD** | **GR** | **95% HPD2** | **FT** | **95% HPD3** | **CW** | **95% HPD4** | **ASD** | **95% HPD5** |
| SM | **0.3896** | **0.224, 0.559** | |  |  |  |  |  |  |  |
| GR | -0.0435 | -0.142, 0.049 | **0.2011** | **0.068, 0.253** | |  |  |  |  |  |
| FT | -0.0088 | -0.050, 0.039 | 0.0206 | -0.008, 0.057 | 0.0472 | 0.018, 0.058 | |  |  |  |
| CW | 0.0339 | -0.058, 0.122 | -0.0028 | -0.067, 0.067 | -0.0105 | -0.051 , 0.012 | 0.1712 | 0.051, 0.224 | |  |
| ASD | 0.0012 | -0.068, 0.074 | -0.0339 | -0.103, -0.001 | -0.0281 | -0.047, 0.001 | -0.0006 | -0.055, 0.046 | 0.1214 | 0.024, 0.119 |
| **Hoffman, NC** | **SM** | **95% HPD** | **GR** | **95% HPD2** | **FT** | **95% HPD3** | **CW** | **95% HPD4** | **ASD** | **95% HPD5** |
| SM | **0.5738** | **0.337, 0.829** | |  |  |  |  |  |  |  |
| GR | 0.0691 | -0.053, 0.209 | **0.2517** | **0.110, 0.352** | |  |  |  |  |  |
| FT | 0.0954 | -0.024, 0.215 | -0.0219 | -0.103, 0.061 | **0.2441** | **0.141, 0.345** | |  |  |  |
| CW | **0.1713** | **0.041, 0.353** | 0.0102 | -0.088, 0.127 | **0.1407** | **0.0351, 0.238** | **0.3582** | **0.171, 0.505** | |  |
| ASD | 0.0669 | -0.051, 0.231 | 0.0138 | -0.086, 0.111 | -0.0149 | -0.103, 0.075 | 0.1148 | 0.016, 0.254 | **0.2879** | **0.128, 0.413** |
| **Ellerbe, NC** | **SM** | **95% HPD** | **GR** | **95% HPD2** | **FT** | **95% HPD3** | **CW** | **95% HPD4** | **ASD** | **95% HPD5** |
| SM | **0.3981** | **0.223, 0.566** | |  |  |  |  |  |  |  |
| GR | -0.0621 | -0.173, 0.049 | **0.2827** | **0.118, 0.375** | |  |  |  |  |  |
| FT | -0.0286 | -0.088, 0.038 | **0.0750** | **0.022, 0.136** | 0.0883 | 0.033 , 0.117 | |  |  |  |
| CW | -0.0102 | -0.094, 0.076 | 0.0085 | -0.063, 0.086 | 0.0255 | -0.021, 0.063 | 0.1582 | 0.033, 0.183 | |  |
| ASD | 0.0161 | -0.069, 0.104 | -0.0158 | -0.092, 0.058 | -0.0390 | -0.075, 0.010 | -0.0124 | -0.077, 0.035 | 0.1633 | 0.022, 0.161 |

**Table S4**: Null **G** matrices estimated for each randomized population along with 95% HPD intervals.

| **Pennsylvania** | **SM** | **95% HPD** | **GR** | **95% HPD** | **FT** | **95% HPD** | **CW** | **95% HPD** | **ASD** | **95% HPD** |
| --- | --- | --- | --- | --- | --- | --- | --- | --- | --- | --- |
| SM | 0.02509 | 0.01, 0.05 |  |  |  |  |  |  |  |  |
| GR | 0.00150 | -0.02, 0.02 | 0.03106 | 0.01, 0.06 |  |  |  |  |  |  |
| FT | -0.00175 | -0.01, 0.01 | -0.00349 | -0.02, 0.01 | 0.01749 | 0.01, 0.03 |  |  |  |  |
| CW | 0.00014 | -0.02, 0.02 | -0.0005 | -0.02, 0.02 | 0.0045 | -0.01, 0.02 | 0.03112 | 0.01, 0.06 |  |  |
| ASD | 0.00081 | -0.02, 0.02 | 0.00104 | -0.02, 0.03 | -0.00054 | -0.02, 0.01 | 0.00188 | -0.02, 0.02 | 0.03215 | 0.01, 0.06 |
| **Maryland** | **SM** | **95% HPD** | **GR** | **95% HPD** | **FT** | **95% HPD** | **CW** | **95% HPD** | **ASD** | **95% HPD** |
| SM | 0.03081 | 0.01, 0.06 |  |  |  |  |  |  |  |  |
| GR | 0.00202 | -0.02, 0.02 | 0.033003 | 0.01, 0.06 |  |  |  |  |  |  |
| FT | -0.00268 | -0.02, 0.01 | -0.00246 | -0.01, 0.01 | 0.01418 | 0.01, 0.02 |  |  |  |  |
| CW | 0.00107 | -0.02, 0.02 | 0.000964 | -0.02, 0.03 | 0.003203 | -0.01, 0.02 | 0.03146 | 0.01, 0.06 |  |  |
| ASD | 0.00076 | -0.02, 0.02 | 0.002035 | -0.03, 0.02 | -0.0033 | -0.02, 0.01 | 0.00226 | -0.02, 0.03 | 0.0287 | 0.01, 0.06 |
| **Hoffman** | **SM** | **95% HPD** | **GR** | **95% HPD** | **FT** | **95% HPD** | **CW** | **95% HPD** | **ASD** | **95% HPD** |
| SM | 0.03046 | 0.01, 0.06 |  |  |  |  |  |  |  |  |
| GR | 0.00222 | -0.02, 0.03 | 0.03103 | 0.01, 0.06 |  |  |  |  |  |  |
| FT | 0.00016 | -0.01, 0.02 | -0.00284 | -0.02, 0.01 | 0.0207 | 0.01, 0.04 |  |  |  |  |
| CW | 0.00253 | -0.02, 0.02 | -0.00055 | -0.03, 0.02 | 0.00704 | -0.01, 0.03 | 0.03543 | 0.01, 0.07 |  |  |
| ASD | 0.00104 | -0.02, 0.0 | 0.00154 | -0.02, 0.03 | -0.00047 | -0.02, 0.02 | 0.00447 | -0.02, 0.03 | 0.03554 | 0.01, 0.07 |
| **Ellerbe** | **SM** | **95% HPD** | **GR** | **95% HPD** | **FT** | **95% HPD** | **CW** | **95% HPD** | **ASD** | **95% HPD** |
| SM | 0.02739 | 0.01, 0.05 |  |  |  |  |  |  |  |  |
| GR | -0.00012 | -0.02, 0.02 | 0.03163 | 0.01, 0.06 |  |  |  |  |  |  |
| FT | -0.00283 | -0.02, 0.01 | -0.00114 | -0.02, 0.02 | 0.01883 | 0.01, 0.03 |  |  |  |  |
| CW | 0.00045 | -0.02, 0.02 | -0.00009 | -0.03, 0.02 | 0.00623 | -0.01, 0.02 | 0.03463 | 0.01, 0.07 |  |  |
| ASD | 0.00107 | -0.02, 0.02 | 0.00228 | -0.02, 0.03 | -0.00213 | -0.02, 0.01 | 0.00169 | -0.02, 0.03 | 0.03531 | 0.01, 0.07 |

**Table S5:** Mean g_max_ of each population. We performed spectral decomposition on each posterior **G** matrix. The first eigenvector of each sample was then used to calculate the mean.

| **Trait** | **Pennsylvania** | **Maryland** | **Hoffman, NC** | **Ellerby, NC** |
| --- | --- | --- | --- | --- |
| Seed mass | 0.9744 | 0.9637 | 0.7741 | 0.8668 |
| Growth rate | -0.2092 | -0.2177 | 0.1120 | -0.4469 |
| Flow. Time | -0.0089 | -0.0433 | 0.2755 | -0.1829 |
| Corolla width | 0.0619 | 0.1441 | 0.5081 | -0.0663 |
| A-S distance | 0.0537 | 0.0340 | 0.2327 | 0.1050 |

**Table S6:** Summary of eigentensor means for observed and random **S** with 95% HPD interval boundaries.

| **Eigentensor** | **Observed** | **Obs. Low** | **Obs. High** | **Random** | **Rnd. Low** | **Rnd. High** |
| --- | --- | --- | --- | --- | --- | --- |
| E1 | 0.09129 | 0.01973 | 0.19793 | 0.00036 | 0 | 0.00122 |
| E2 | 0.01680 | 0.00057 | 0.03943 | 3.00E-04 | 0 | 0.00092 |
| E3 | 0.00828 | 4.00E-05 | 0.02308 | 0.00027 | 0 | 0.00074 |

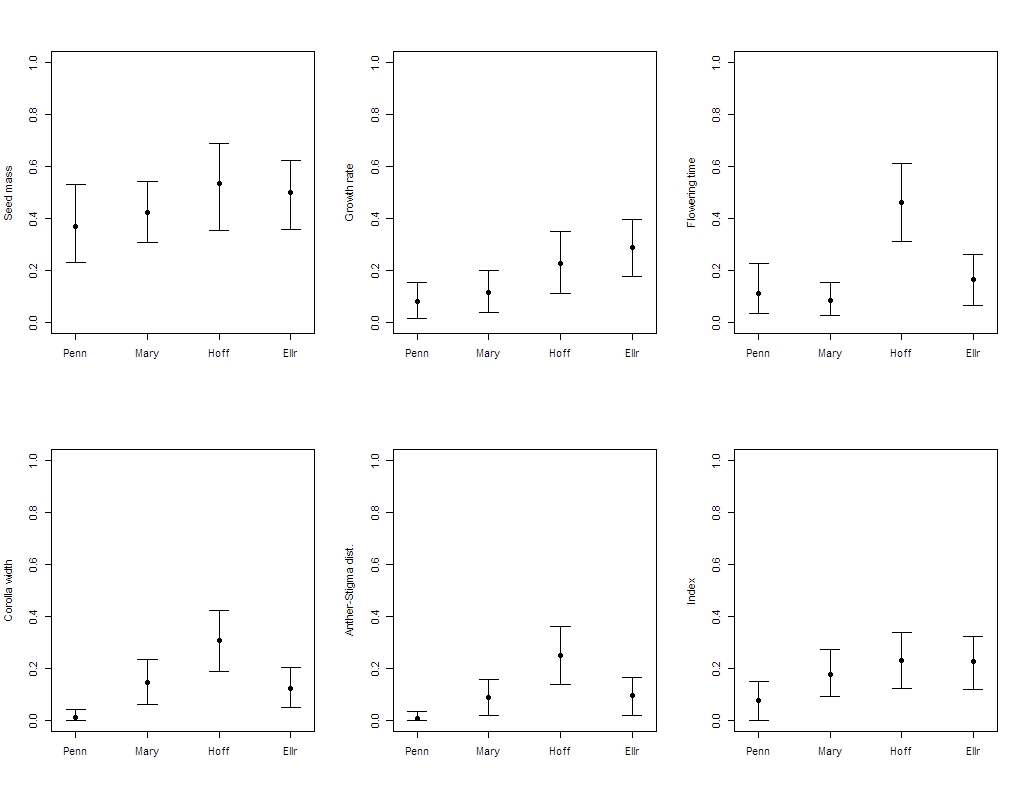

**Figure S1:** Broad sense heritability estimates for each trait and population, estimated using univariate Bayesian generalized linear mixed-effect models.

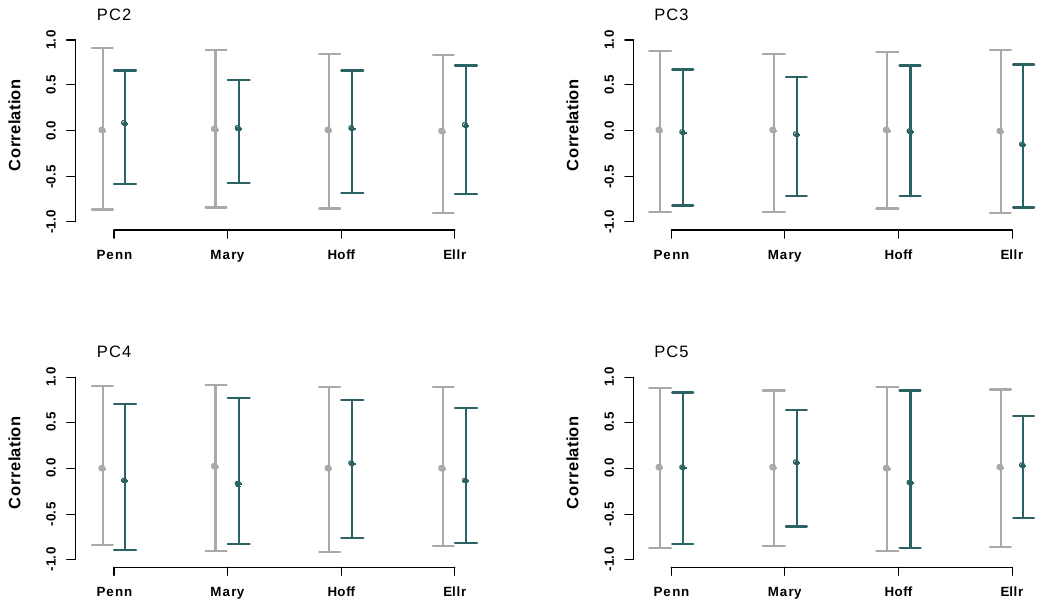

**Figure S2:** The correlation between Principal Components 2 through 5 of each population **G** matrix and the vector of greatest multivariate clinal divergence (Observed, in blue) compared to the correlation between the g_max_ of each population **G** matrices and randomized vectors (Randomized, in gray). Error bars represent the 95% HPD intervals. “Penn” is Pennsylvania, “Mary” is Maryland, “Hoff” is Hoffman, NC and “Ellr” is Ellerby, NC.

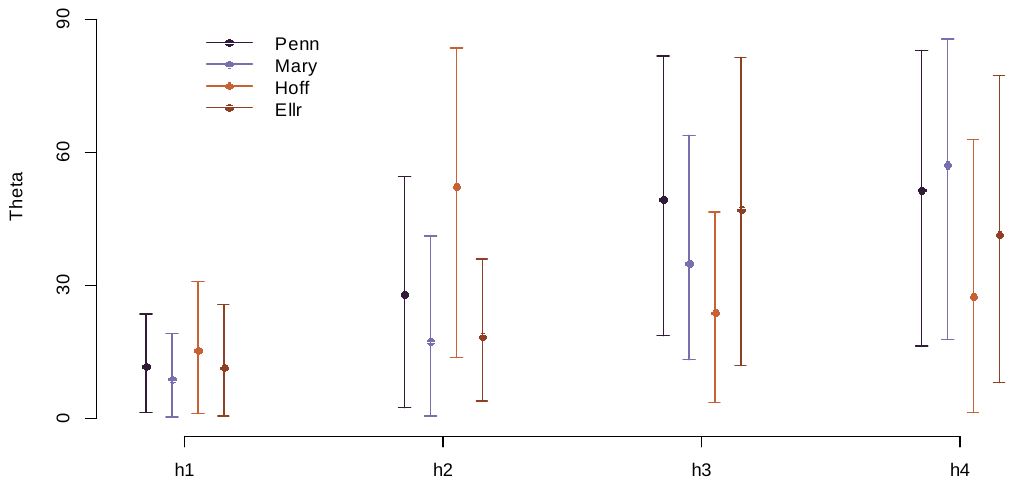

**Figure S3:** Angle in degrees between populations and the shared subspace **H**. The angles are typically small, with large, overlapping HPD intervals, meaning differences between the populations and the shared trait subspace are minimal.

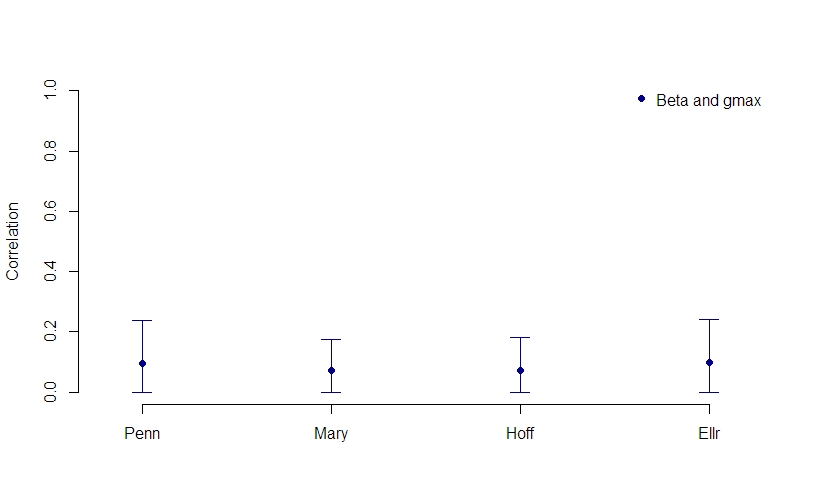

**Figure S4:** The correlation between the estimated selection gradient, β, and the first eigenvector of the **G** matrix, g_max_, across all posterior samples for each population. Betas were calculated by taking the inverse of the population **G**s and multiplying by the variance standardized axis of greatest multivariate divergence, Cline_max_.
